# Connectivity and dispersal mode shape the landscape genetics of a carnivorous pitcher plant-arthropod metacommunity

**DOI:** 10.64898/2026.08.09.743144

**Authors:** Nonno Hasegawa, Asa E. Conover, Matin Miryeganeh, David W. Armitage

**Author notes:** These authors contributed equally (Nonno Hasegawa, David W. Armitage).

## Abstract

Dispersal differences between hosts and their symbionts can generate mismatched population structure, potentially destabilizing beneficial interactions across space. We tested this possibility in the carnivorous pitcher plant *Darlingtonia californica* and its obligate arthropod associates, the midge *Metriocnemus edwardsi* and the mite *Sarraceniopus darlingtoniae*, sampled across sites spanning the host’s patchy range in Oregon and northern California, USA. Comparing nuclear and chloroplast genomic data from *D. californica* with mitochondrial COI data from both arthropods, we tested how range position, landscape connectivity, and dispersal mode influence population genetic structure across this mutualistic metacommunity. Host plant populations supported the central–marginal hypothesis: nuclear diversity declined toward the range margins, and marginal populations showed greater nuclear genetic differentiation. Chloroplast variation was more weakly structured, most clearly separating the northern Oregon Coast populations and revealing cytonuclear discordance consistent with historical seed-mediated movement or chloroplast capture near the boundary between neighboring regions. Landscape connectivity estimated from an ecological niche model was also associated with genetic exchange. Circuit-theoretic current flow was positively related to effective migration inferred independently from plant genotypes. Further, landscape resistance explained variation in plant and mite differentiation beyond geographic distance alone. Both arthropods showed significant spatial congruence with the host plant but not with one another, a pattern inconsistent with co-dispersal and suggesting that each associate tracks the shared landscape according to its own dispersal biology. These results show that regional genetic concordance among obligate ecological partners can coexist with substantial differences in the processes governing their movement and local connectivity.

## Introduction

Species with patchy or fragmented distributions are anticipated to experience reduced connectivity among populations, leading to stronger genetic drift, lower within-population diversity, and increased genetic differentiation across landscapes (Cheptou et al. 2017; Schlaepfer et al. 2018). These effects may be more pronounced near range margins, where populations often occupy smaller, more isolated habitat patches and experience more environmental stress relative to those near the range center. This expectation forms the basis of the *central–marginal hypothesis* (CMH), which predicts that peripheral populations should generally harbor lower genetic diversity and are more strongly differentiated than central populations (Villellas et al. 2014; Langin et al. 2017; Kennedy et al. 2020). However, empirical support for this pattern remains mixed, particularly in patchy habitat-specialist species where historical range dynamics, dispersal ability, and landscape structure may obscure simple core–periphery gradients.

Landscape genetics provides a framework for linking these spatial genetic patterns to the environmental and geographic features that shape gene flow (Wang, Savage, et al. 2009; Epps et al. 2007; Balkenhol et al. 2015). In particular, ecological niche models can be used to derive landscape resistance surfaces, which can then be combined with circuit-theoretic connectivity models to predict movement among habitat patches (McRae et al. 2016). Comparing these predictions with empirically inferred gene flow from genomic data provides a direct test of whether independently modeled habitat connectivity explains observed genetic exchange (Marcus et al. 2021). Such approaches are especially valuable for rare or threatened species, where future habitat change may further disrupt connectivity and increase extinction risk in already fragmented systems (Hanski et al. 2017; Cheptou et al. 2017).

Comparative landscape genetic approaches can also reveal whether co-distributed interacting species respond similarly to their shared landscape. In obligate symbioses, partners may share habitats but markedly differ in dispersal mode, potentially producing discordant genetic structure even when their ecological distributions are tightly linked (Brucker and Bordenstein 2012). Conversely, shared dispersal routes or co-dispersal (e.g., phoretic) associations may generate concordant genetic structure among partners. Testing these alternatives can help link alleles’ movement across fragmented landscapes with how species interactions are maintained over evolutionary time (Bohonak 1999). However, few studies have simultaneously examined the host and multiple obligate symbionts within a single landscape genetic framework.

Carnivorous pitcher plants provide a useful system for such analyses because they are restricted to nutrient-poor, spatially discrete habitats and support specialized arthropod communities in their fluid-filled leaves. *Darlingtonia californica* Torr., the ’cobra lily’, is the sole species in its genus and is endemic to serpentine seeps and fens in coastal Oregon and northern California, USA (**Fig. 1A**). Its strict hydrological requirement for cold, flowing water (Adlassnig et al. 2005) and preference for serpentine soils makes suitable habitat extremely rare and very patchily distributed across its topographically complex landscape. This geography makes *D. californica* an ideal system for investigating how range position and landscape connectivity predict genomic diversity and gene flow.

**Figure 1:**
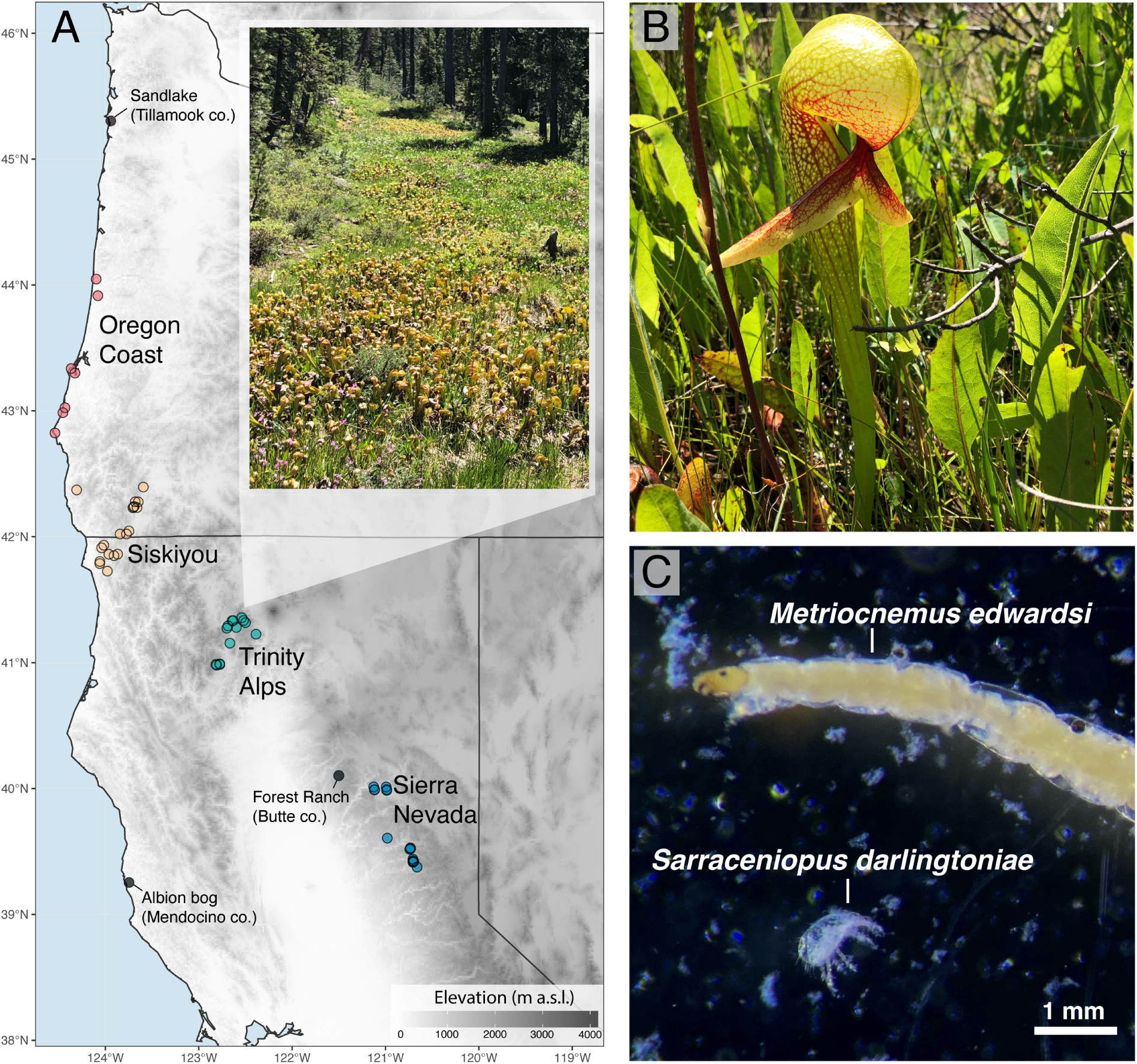
**(A)** Map of sampling points (*n* = 59) in N. California and Oregon, USA, colored by geographic region. Labeled grey points show putative introduced populations. Inset depicts a typical montane fen containing a large *Darlingtonia californica* population. **(B)** Individual pitcher leaf of *D. californica*. **(C)** Midge and mite symbionts *Metriocnemus edwardsi* and *Sarraceniopus darlingtoniae* found in the pitcher’s digestive zone. Photos by DWA and NH.

The pitchers of *D. californica* (**Fig. 1B**) also contain obligate arthropod symbionts that contribute to prey breakdown and nutrient cycling. Two such species are the detritivorous midge *Metriocnemus edwardsi* Jones 1916 and the mite *Sarraceniopus darlingtoniae* Fashing & O’Connor 1984 (Naeem 1988; Nielsen 1990; Fashing and O’Connor 1984; Fashing 2004) (**Fig. 1C**). These species differ strongly in their dispersal traits. Adult *M. edwardsi* are winged and capable of active dispersal, whereas *S. darlingtoniae* disperses locally as a deutonymph by walking among nearby pitchers. Long-distance dispersal by the mite remains unresolved, although laboratory observations suggest that deutonymphs can attach to adult midges, raising the possibility of phoretic dispersal (Fashing 2004). If mites disperse via midges, the two arthropods should share more genetic structure than either does with the host plant. The mechanisms of pollen and seed dispersal in *D. californica* are also poorly understood, although bee pollination and zoochorous seed dispersal have been proposed (Meindl and Mesler 2011; Collingsworth 2015). Because nuclear loci are transmitted by both pollen and seed whereas the chloroplast genome is maternally inherited and dispersed by seed alone, comparing differentiation between the two genomes can provide information on the dispersal biology of this poorly characterized species.

Here we present a range-wide landscape genetic analysis of *D. californica* and two obligate arthropod symbionts across the host plant’s geographic range. Combining nuclear and chloroplast SNPs derived from double-digest restriction-site associated DNA sequencing (ddRADseq) of the host plant with mitochondrial COI sequence data from both arthropods, we ask: (i) whether each taxon shows population structure consistent with the host plant’s patchy distribution, and whether marginal populations show the reduced diversity and elevated differentiation predicted by the central–marginal hypothesis; (ii) whether landscape connectivity derived from niche-model resistance surfaces predicts effective migration estimated independently from the genomic data; (iii) whether isolation by distance and isolation by resistance contribute differently to genetic structure across the three taxa; and (iv) whether the arthropods’ spatial genetic structure is congruent with that of their host, or whether each tracks the landscape according to its own dispersal biology.

## Materials and Methods

### Sample collection

Between May and July 2022, we collected tissues from juvenile *Darlingtonia* leaves across 59 sites spanning the species’ entire geographic range across Oregon and California, USA (**Fig. 1A**). Within this range, replicate sites were visited within all discrete regions in which they are known to occur, including all sites near the species’ latitudinal range margins. We also sampled introduced *Darlingtonia* populations in Albion Bog, CA (Mendocino Co.), Forest Ranch, CA (Butte Co.), and a site of unknown status in Sandlake, OR (Tillamook Co.). Population sizes at each site varied from 3 to over 10,000 individual plants. From up to eight different plant ramets at each site, we collected a single, developing (unopened) pitcher leaf and immediately preserved it in silica gel. We additionally collected mites and midge larvae from an arbitrary number of individual mature plants by dissecting the base of a pitcher leaf from the plant. Immediately following collection, mites and midges were recovered under a microscope, concentrated into vials containing a DNA/RNA protection reagent (New England Biolabs, Inc., Ipswich, MA) and stored at 4°C. Permits for collections at all sites are available on request.

### Plant sequence processing

Full DNA extraction, library preparation, sequencing, and organelle-reference assembly protocols are provided in the Supplementary Methods. Briefly, double-digest restriction-site associated DNA sequencing (ddRADseq) libraries were generated for 199 *Darlingtonia* samples, with at least three individuals represented per collection site.

Demultiplexed ddRADseq reads were processed for quality and trimmed to 148 bp using the process_radtags pipeline in Stacks v2.58 (Rochette et al. 2019) with default options. Reads were first mapped to the assembled chloroplast genome using Bowtie2 (Langmead and Salzberg 2012), and mapped reads were retained as chloroplast DNA (cpDNA). Unmapped reads were then aligned to the assembled mitochondrial genome, and reads remaining unmapped after this step were treated as predominantly nuclear DNA (nuDNA). Nuclear reads were processed using the denovo_map.pl pipeline, which performs *de novo* assembly of restriction-site flanking regions, phasing, SNP calling, and alignment, with the parameter settings -M 3 -n 3 --rm-pcr-duplicates.

Because the chloroplast genome is haploid, cpDNA variants were called directly from the Bowtie2 alignments using bcftools (Danecek et al. 2021) under a haploid model (--ploidy 1). We retained alignments with mapping quality ≥ 30, biallelic SNPs with site quality ≥ 30 and total depth ≥ 10, and set genotypes supported by fewer than two reads to missing. Sites genotyped in at least 30% of individuals were retained, yielding 42 high-confidence plastid SNPs. Because per-locus plastid coverage from ddRADseq was low (mean ∼ 4×), chloroplast data were only used to explore maternal-lineage structure qualitatively through principal component analysis and not used for any statistical inference.

### Arthropod sequence processing

Full DNA extraction, COI amplification, and sequencing protocols are provided in the Supplementary Methods. Site-level pooled samples were successfully sequenced for 57 midge and 45 mite collections.

Sequences were trimmed by 5 bp from the start and 20 bp from the end, and bases with quality scores below Q20 were removed using seqtk. Forward and reverse reads were joined using fastq_join in USEARCH v12. Because the approximately 700-bp COI amplicon exceeded the maximum joined length obtainable from the 600-cycle reads, reads were joined across the intervening gap before dereplication and chimera filtering in USEARCH and *de novo* OTU clustering with Swarm v3 (Mahé et al. 2021). Clusters were ranked by read count, and representative sequences were assigned taxonomically using BLAST searches against the NCBI non-redundant nucleotide database. Clusters returning Chironomidae or Sarcoptiformes matches were retained.

To verify taxonomic placement and identify clusters most closely matching the target taxa, we constructed COI phylogenies from sequences available in NCBI, including *Metriocnemus* midges and Histiostomatidae mites. Trees were inferred using FastTree2 with default parameters and used to exclude clusters falling outside the target midge genus or mite family. The highest-abundance target cluster from each site was retained for population genetic analysis. Midge clusters were mapped to the NCBI *Metriocnemus* reference sequence HQ551710.1 using BWA-MEM, and variants were called using bcftools. The resulting VCF was analyzed using the populations module in Stacks to obtain population genetic summary statistics. Because no external *Sarraceniopus* COI reference was available, a high-quality mite cluster from Cape Blanco, Oregon was used as the reference.

### Population genetic metrics

For *D. californica* nuclear and chloroplast DNA datasets and for COI datasets of both arthropod species, we calculated population genetic metrics at multiple spatial scales, including individual leaf samples (*n* = 199; plant samples only), sampling sites (*n* ≤ 46, depending on sample type), and geographic clusters of sites (*n* = 4; Oregon Coast, Siskiyou, Trinity, and Sierra Nevada). For each dataset and spatial scale, genetic diversity within sampling units was quantified using nucleotide diversity (*π*) and, where appropriate, observed (*H_O_*) and expected heterozygosity (*H_E_*) and inbreeding coefficients (*F_IS_*), with heterozygosity- and inbreeding-based metrics calculated only for diploid nuclear loci. Pairwise fixation indices (*F_ST_* ; Weir and Cockerham 1984) were calculated among sampling units using the R package hierfstat (R Core Team 2024; Goudet 2005). To evaluate the spatial scale of nuclear population genetic structure in *D. californica*, we conducted a hierarchical analysis of molecular variance (AMOVA) using individual-level genotypes from pruned nuclear SNPs. Individuals were nested within sampling sites, and sampling sites were nested within four geographic regions: Oregon Coast, Siskiyou, Trinity, and Sierra Nevada, excluding introduced populations. Pairwise genetic distances among individuals were calculated from the SNP data and partitioned into variance components comprising regions, sites within regions, individuals within sites, and variation within individuals. Significance of each variance component was assessed using 999 restricted permutations implemented in the poppr (Kamvar, Tabima, et al. 2014; Kamvar, Brooks, et al. 2015) and ade4 (Dray and Dufour 2007) R packages. Because ddRADseq samples a restricted subset of each genome, these estimates represent diversity across the recovered ddRAD loci rather than genome-wide nucleotide diversity.

### Population structure

Population structure in the *D. californica* nuclear SNP dataset was further evaluated using principal component analysis (PCA) and ancestry inference. To construct a set of approximately unlinked SNPs, variants were filtered in PLINK v1.9 (Chang et al. 2015) using a minor allele frequency threshold of 0.01, a maximum locus missingness threshold of 90%, and a maximum individual missingness threshold of 95%. Linkage disequilibrium pruning was then performed using a sliding window of 50 SNPs, a step size of 10 SNPs, and an *r*^2^ threshold of 0.2, resulting in a final dataset of 19,166 SNPs across 199 individuals. PCA was conducted in PLINK, and individual ancestry proportions were estimated using ADAMIXTURE (Saurina-i-Ricos et al. 2026) for *K* = 2 − 15, with the optimal range of *K* assessed using five-fold cross-validation (CV). Cross-validation supported *K* = 3 − 4 as the most plausible range of ancestry clusters. Following robustness recommendations (Hemstrom et al. 2024), we repeated ancestry analyses across minor allele frequency thresholds of 0.01 and 0.05 and locus missingness thresholds of 5%, 10%, 20%, 50%, 70%, and 90%. These sensitivity analyses frequently supported *K* = 4, particularly under stricter missingness filters, suggesting that the fourth ancestry component was not an artifact of the relaxed filtering used in the primary analysis. We therefore interpret *K* = 3 as representing the most parsimonious level of nuclear population structure and *K* = 4 as a plausible finer-scale partition. To assess whether additional structure was present within these groups, we repeated ancestry analyses within each of the four major *K* = 4 clusters. These analyses provided no evidence for further subdivision (Supplementary Table S3).

### Testing the central–marginal hypothesis

We tested two predictions of the central–marginal hypothesis: that populations toward the geographic margins of the species’ range should have lower genetic diversity and greater genetic isolation than populations nearer the range center. We defined the geographic center of the sampled range as the point corresponding to the median latitude and median longitude of our collected *Darlingtonia* occurrences and calculated the Haversine distance between each site and this centroid. To test whether genetic diversity declined toward the range margins, we restricted the diversity analysis to the plant nuclear dataset, for which diversity estimates were based on individually resolved genotypes. The chloroplast dataset was excluded because its low per-locus ddRAD coverage precluded reliable site-level diversity estimation, and the mite and midge DNA was extracted from pooled samples containing multiple individuals, so conventional within-population nucleotide diversity could not be calculated.

For the nuclear dataset, we modeled nucleotide diversity (*π*) as a function of distance from the range centroid and estimated population size, which was coded as a categorical variable. Populations were classified as small when they contained fewer than 100 plants and as large when they contained many hundreds to thousands of plants. Linear regression models were fitted with distance standardized to have a mean of zero and a standard deviation of one. Because the central–marginal hypothesis predicts that nucleotide diversity declines with increasing distance from the range center, we evaluated the distance coefficient using a one-sided hypothesis test with the alternative hypothesis *β*_distance_ *<* 0. We similarly used a one-sided test for the population-size effect, with the alternative hypothesis that small populations have lower nucleotide diversity than large populations.

To test whether populations at the range margins were more genetically isolated, we classified the Oregon Coast and Sierra Nevada regions as marginal and the Siskiyou and Trinity regions as central. For each plant and arthropod genetic dataset, we calculated a population-level measure of genetic differentiation by averaging all pairwise *F_ST_* values between a focal population and the other sampled populations within its original geographic region. Only within-region population pairs were included so comparisons between (for example) Oregon Coast and Sierra Nevada populations were excluded despite sharing the same marginal classification. We then compared population-level mean *F_ST_* between central and marginal populations using linear regression, with central populations treated as the reference group. A positive marginal-population coefficient, corresponding to higher mean within-region *F_ST_* at the range margins, was interpreted as support for the prediction of greater marginal genetic isolation.

### Landscape resistance and connectivity

We used ecological niche modeling (ENM) to approximate landscape resistance for *Darlingtonia californica*. We gathered georeferenced records (*n* = 169) of *Darlingtonia* patches from our field observation data, which was itself concatenated from records shared on GBIF and through personal communications with users of the community science platform iNaturalist (www.inaturalist.org). All analyses were conducted at a spatial resolution of 0.5 arcmin (approximately 66 ha at 40°N) across a study region spanning the Northwestern United States.

Candidate environmental predictors comprised the 19 bioclimatic variables from the WorldClim 2 database (Fick and Hijmans 2017); seven soil properties at 15 cm depth from SoilGrids (Poggio et al. 2021); four consensus land-cover classes (Tuanmu and Jetz 2014); Euclidean distance to the nearest mapped spring or seep; and a binary raster of ultramafic (serpentine) substrate derived by rasterizing mapped ultramafic outcrop polygons (Garnica-Díaz et al. 2023) (all covariates are listed in Supplementary Table S1). The calibration area was defined as the union of 150 km buffers placed around each occurrence record, restricting model fitting to the region accessible to the species. Within this extent we generated 10,000 random background points. Collinearity among the 31 continuous predictors was reduced using a hierarchical filtering algorithm (Adde et al. 2023), which identifies pairs of predictors with |*ρ*| *>* 0.8 and retains the variable with higher predictive performance.

We fit MaxEnt models (Phillips, Anderson, and Schapire 2006; Phillips, Anderson, Dudík, et al. 2017) to the retained predictors using the ENMeval 2.0 R package (Kass et al. 2021), tuning across three feature-class combinations (linear; linear–quadratic; linear–quadratic–hinge) and eight regularization multipliers (1.5–5.0 in increments of 0.5), for 24 candidate models. Model performance was evaluated using 4-fold spatial block cross-validation. We selected the model maximizing the mean Continuous Boyce Index (Hirzel et al. 2006) and generated cloglog suitability predictions across the study extent.

We transformed this suitability raster (on the [0, 1] scale) to a resistance raster (on the [1, 1000] scale) using the linear function *R* = 1000 − 999*S* where *S* is a grid cell’s suitability and *R* is its resistance. This assumes that site suitability is linearly proportional to its ability to spread across the landscape, which is an appropriate null expectation for sessile habitat specialists (Keeley et al. 2016). We further truncated all cells falling above a suitability threshold *τ* to *R* = 1, defined as the minimum predicted suitability value that correctly predicts the presences of at least 66% of our occurrence records. Geographic and resistance distances were then calculated between plant sampling sites using the R package gdistance (Etten 2017) using the Haversine formula or the commute distance algorithm (Fouss et al. 2007), respectively.

We additionally used Omniscape.jl ver. 0.6.2 (Landau et al. 2021) to estimate regional landscape connectivity from the ENM-derived resistance surface. Omniscape (McRae et al. 2016) is a circuit-theoretic connectivity algorithm that moves a circular window of radius *r* across the landscape and, for each focal pixel, treats all habitat pixels within the window as current sources and the central pixel as the target. We used a window radius of 1000 pixels and defined core habitat as pixels with resistance values of 1. Accumulated current flow was summed across all window positions, producing a continuous surface in which pixel values reflect the degree to which a location is expected to facilitate movement under a random-walk dispersal model.

### Modeling isolation-by-distance and resistance

We jointly assessed isolation-by-distance (IBD) and isolation-by-resistance (IBR) using a Bayesian analogue of the maximum-likelihood population-effects (MLPE) model to account for the non-independence of pairwise genetic distances (Clarke et al. 2002). Because pairwise geographic and resistance distances were strongly correlated (Mantel *r* = 0.93), we first regressed resistance distance on geographic distance and used the resulting residuals (*resid resist*) as an orthogonalized predictor (Supplementary Figure S1). This parameterization isolates the contribution of landscape resistance independent of geographic distance, allowing simultaneous estimation of both effects on linearized genetic differentiation (*F_ST_ /*[1 − *F_ST_*]). Both predictors were standardized to zero mean and unit variance so that slopes are comparable among taxa and marker types.

Models were specified with fixed effects for geographic distance and residualized resistance distance, and a random intercept over the two populations contributing to each comparison. This structure models covariance among *F_ST_* values that share a population, reproducing the MLPE error structure. Models were fitted in R (v4.4) using brms (Bürkner 2017), an interface to the programming language Stan (Carpenter et al. 2017) using default uninformative priors. Each model was run with six chains of 10,000 iterations (5,000 warmup), yielding 30,000 post-warmup draws. Convergence was verified using *R^*, bulk and tail effective sample sizes (Vehtari et al. 2017), and graphical posterior predictive checks. This procedure was applied separately to pairwise *F_ST_* matrices derived from *Darlingtonia* nuclear SNPs and from *Metriocnemus* and *Sarraceniopus* COI sequences. Support for each effect was summarized as the proportion of posterior draws greater than zero (*P* [*β >* 0]), together with posterior medians and 95% credible intervals.

### Inference of effective migration surfaces

To visualize spatial variation in gene flow across the range of *D. californica*, we estimated fast effective migration surfaces (fEEMS) from the nuDNA dataset (Marcus et al. 2021). We used the fEEMS 2.0 Python package to estimate spatial patterns of effective migration by modeling genetic dissimilarity among sampling locations. Input data consisted of nuclear SNP-based genotypes and geographic coordinates for each sampling site. Following the software authors’ implementation, we fixed the *α* parameter to the inverse of the estimated constant edge weight and used a leave-one-out cross-validation procedure to find the value of smoothing parameter *λ* that minimized the mean squared error between predicted and observed allele frequencies (**Supplementary Figure S2**). The fitting procedure then output a spatial grid where edge weights represent the posterior mean for effective migration between nodes connecting them. This mesh was then converted to a spatial raster.

### Comparing migration and connectivity surfaces

We compared spatial patterns of inferred gene flow with independently derived landscape connectivity estimates using effective migration rates inferred with fEEMS and cumulative current flow predicted by Omniscape. The two raster surfaces were first resampled to a common extent and aggregated by a factor of 10 to produce a shared spatial resolution. This aggregation reduced fine-scale spatial variation and minimized gaps associated with the mesh-based fEEMS estimates. Values from both rasters were then extracted for the same set of 870 grid cells, together with the geographic coordinates of each cell. We then tested whether spatial variation in effective migration was associated with landscape connectivity after accounting for shared geographic structure using a generalized additive model (GAM). Effective migration rate inferred using fEEMS was treated as the response variable, and Omniscape cumulative current flow was included as the predictor of interest after transformation as log_10_(1 + current flow). Spatial autocorrelation was represented using a two-dimensional Gaussian-process smooth of latitude and longitude with a Matérn-type covariance structure:

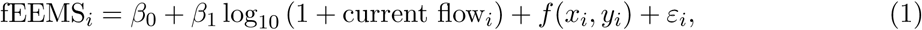

where *f* (*x_i_, y_i_*) represents spatial variation not attributable to connectivity and *ε_i_* is the residual error term. Models were fit using restricted maximum likelihood (REML) in the R package mgcv (Wood 2011). We compared this model with a spatial-only null model. Adding connectivity significantly improved model fit relative to the spatial-only model. For visualization, a partial regression plot was generated using the R package visreg (Breheny and Burchett 2017), showing the estimated effect of connectivity on effective migration while holding the spatial smooth constant.

### Population genetic congruence across species

To assess whether geographic population genetic structure was congruent across *D. californica* and its associated arthropods (*M. edwardsi* and *S. darlingtoniae*), we performed pairwise and three-way generalized Procrustes analyses (GPA) of principal coordinate analysis (PCoA) scores derived from pairwise *F_ST_*matrices. This approach evaluates whether the relative arrangement of sampled localities in genetic-distance space is congruent among organisms after removing arbitrary differences in translation, scale, rotation, and reflection among ordination configurations (**Supplementary Figure S3**).

For each organism, we performed PCoA on its *F_ST_*matrix using the ape R package (Paradis and Schliep 2019), with a Cailliez correction for non-Euclidean distances. The first two PCoA axes were retained, centered, and scaled to unit centroid size before alignment. Pairwise Procrustes analyses were performed for each species pair. For each comparison, the two ordination configurations were aligned, and their congruence was summarized using the Procrustes correlation, with values approaching one indicating greater correspondence between configurations. We also report Mantel correlations between the corresponding pairs of *F_ST_* matrices for comparability with earlier comparative phylogeographic studies, while noting that Procrustean superimposition is more powerful than matrix correlation for this class of problem (Peres-Neto and Jackson 2001).

For the three-way analysis, the nuclear plant, midge, and mite configurations were aligned using an iterative GPA algorithm that minimized disagreement with a consensus configuration (Gower 1975). Congruence was quantified as the mean squared distance of each organism-specific score from the centroid of the corresponding locality across species:

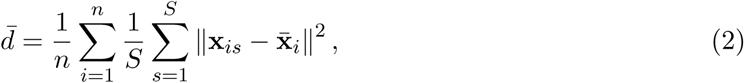

where (*n*) is the number of localities shared among all datasets, (*S* = 3) is the number of species, (**x***_is_*) is the aligned coordinate vector for locality (i) in species (s), and (**x̄***_i_*) is the centroid of locality (i) across species. Smaller values of (*d̂*) indicate stronger congruence, with perfect correspondence across all species and sites when *d̂* = 0.

Statistical significance was assessed using permutation tests with *n*_perm_ = 9999. For each pairwise analysis, locality labels in one configuration were randomly reassigned while the other configuration was held fixed, the Procrustes alignment was recomputed, and the observed Procrustes correlation was compared with the resulting null distribution. Pairwise *P* -values were calculated as the proportion of permutations producing a Procrustes correlation at least as large as the observed value. For the three-way analysis, locality labels were randomly reassigned in one or more focal datasets, the GPA was recomputed, and the resulting mean squared centroid deviation was compared with the observed value. We considered four permutation scenarios: (*i*) midge labels permuted while the plant and mite configurations were held fixed; (*ii*) mite labels permuted while the plant and midge configurations were held fixed; (*iii*) nuclear plant labels permuted while both arthropod configurations were held fixed; and (*iv*) midge and mite labels permuted independently while the plant configuration was held fixed. Because smaller values of (*d̂*) indicate stronger three-way congruence, *P* -values were calculated from the lower tail of the null distribution as the proportion of permutations producing a mean squared centroid deviation less than or equal to the observed value. The three-way analysis included (*n* = 15) localities shared among all three datasets.

## Results

### *Darlingtonia* population structure

*Darlingtonia* sequencing yielded 241,611 nuclear SNPs. Because the chloroplast genome is haploid, plastid variants were called separately under a haploid model, yielding only 42 high-confidence chloroplast SNPs because plastid reads made up only a small share of the ddRAD data. After filtering and linkage pruning for ancestry analysis, the nuclear dataset comprised 19,166 SNPs across 199 individuals. COI barcoding yielded site-level data for 38 *M. edwardsi* (midge) and 19 *S. darlingtoniae* (mite) populations.

Ancestry inference from *Darlingtonia* nuclear SNPs was consistent with the geographic clustering of sites across the species’ range (**Fig. 2A**). Across 12 filtering schemes spanning minor allele frequency and missingness thresholds, cross-validation error was minimized at *K* = 4 in eight cases, *K* = 3 in three, and *K* = 5 in one, while *K* = 2 and *K* ≥ 6 were never favored (**Supplementary Table S2**). We therefore treat *K* = 4 as the best-supported partition, corresponding to the four sampled geographic regions, with *K* = 3 representing a coarser alternative. At *K* = 5, Siskiyou samples separated into northern and southern groups, hinting at unresolved substructure within the largest and most densely sampled region, yet neither this nor any other substructure was supported when each region was analyzed separately (**Supplementary Table S3**). The introduced Albion Bog and Forest Ranch populations clustered with Sierra Nevada sites, consistent with a Sierra Nevada origin for these introductions, and the Sandlake, OR samples clustered within the Oregon Coast population.

**Figure 2:**
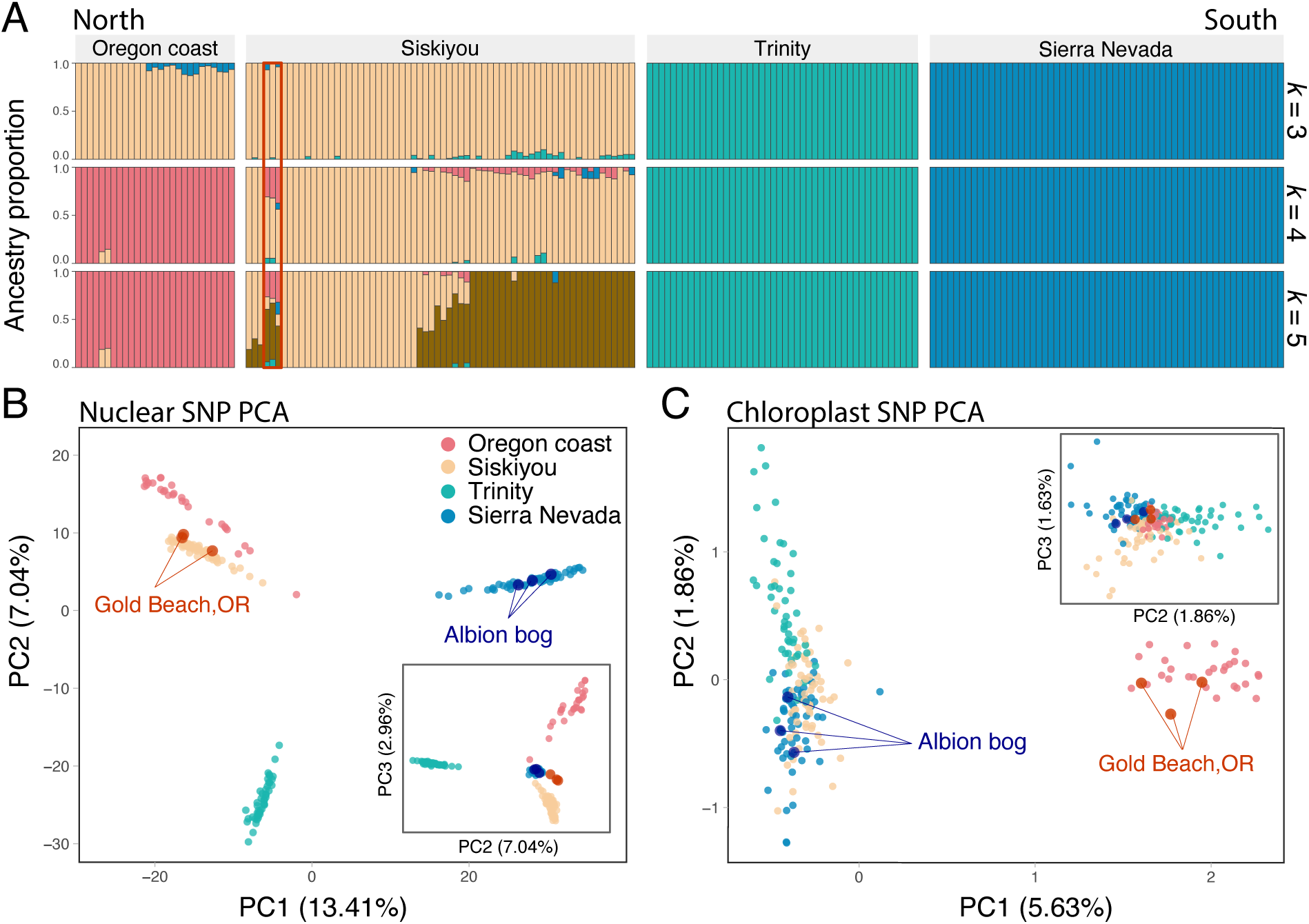
**(A)** ADAMIXTURE-predicted population structure for *D. californica* using nuclear SNP data across three values of *K*. The red box indicates the Gold Beach, OR samples. **(B)** Principal component plot of population genetic differentiation among *D. californica* nuclear DNA samples. **(C)** Principal component plot of population genetic differentiation among *D. californica* chloroplast DNA samples. Inset graphs show additional PC axes. Points are colored by collection region.

Principal component analysis recovered the four geographic regions as largely distinct clusters, with the first three axes explaining 23.4% of genetic variance (**Fig. 2B**). Chloroplast structure was both weaker and differently organized: PC1 clearly separated Oregon Coast samples, while the Siskiyou, Trinity, and Sierra Nevada regions were largely indistinguishable from one another (**Fig. 2C**). All three Gold Beach plant specimens fell within the Siskiyou nuclear cluster yet carried the Oregon Coast–type plastid signature, matching Oregon Coast reference samples at a higher proportion of shared sites than they matched other Siskiyou samples (0.84–0.89 versus 0.58–0.79).

Hierarchical AMOVA confirmed that nuclear SNP variation was structured at multiple spatial scales (**Table 1**). Region accounted for 27.4% of total variation and sites within regions a further 16.8%, with all components significant (*P* = 0.001). Variation among individuals within sites accounted for the largest share (51.6%), while within-individual variation accounted for only 4.2%, a heterozygote deficit consistent with the gap between observed and expected heterozygosity in all regions (**Table 2**). Every sampled individual carried a unique multilocus genotype, and the smallest pairwise distance between any two individuals in the dataset was 2.7% of scored loci, confirming that the leaves we sampled represent distinct genets rather than clonal replicates.

**Table 1:**
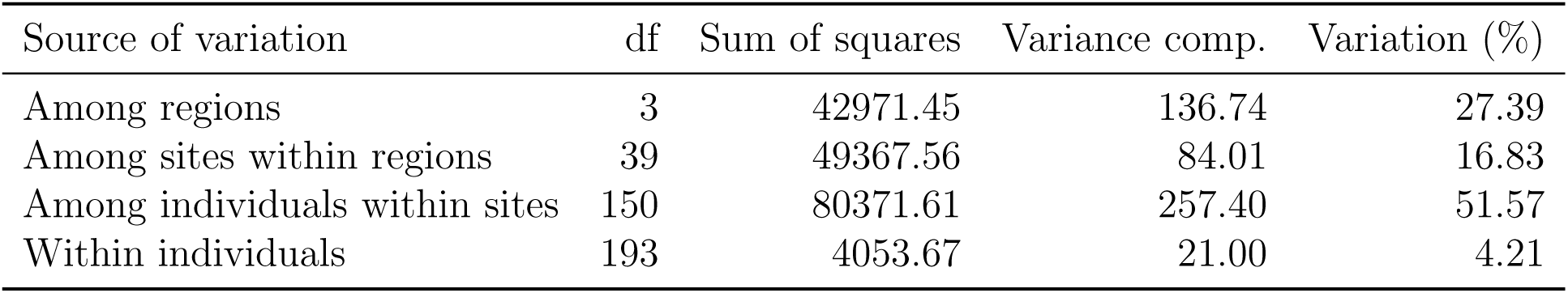
Hierarchical analysis of molecular variance (AMOVA) for *D. californica* nuclear SNP data. Genetic variation was partitioned among geographic regions, among sampling sites within regions, among individuals within sites, and within individuals. The listed variance components were significant at (*P* = 0.001).

**Table 2:**
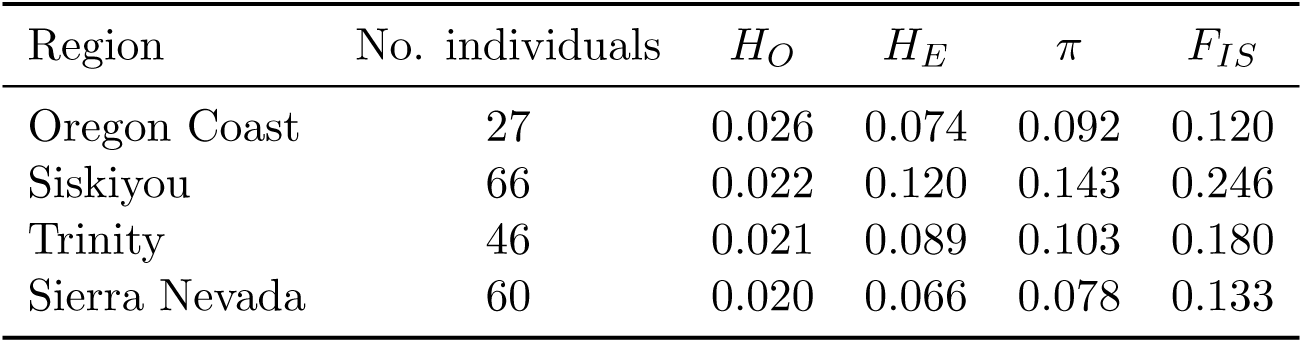
*Darlingtonia californica* population genetic metrics for the nuclear genome, presented at the region scale. Observed heterozygosity (*H_O_*), expected heterozygosity (*H_E_*), nucleotide diversity (*π*), and the inbreeding coefficient (*F_IS_*) are averaged across the sites sampled within each region. Introduced populations are excluded.

| Region | No. individuals | $H_O$ | $H_E$ | $\pi$ | $F_{IS}$ |
| --- | --- | --- | --- | --- | --- |
| Oregon Coast | 27 | 0.026 | 0.074 | 0.092 | 0.120 |
| Siskiyou | 66 | 0.022 | 0.120 | 0.143 | 0.246 |
| Trinity | 46 | 0.021 | 0.089 | 0.103 | 0.180 |
| Sierra Nevada | 60 | 0.020 | 0.066 | 0.078 | 0.133 |

### Central–marginal effects

Nuclear nucleotide diversity (*π*) declined strongly with increasing distance from the range centroid after accounting for population size (*β* = −0.022, *t*_40_ = −6.87, one-sided *P <* 0.0001; **Fig. 3A**). Population size also had a strong effect, with small populations containing lower nuclear diversity than large populations at a given distance from the range centroid (*β* = −0.034, *t*_40_ = −5.30, one-sided *P <* 0.0001).

**Figure 3:**
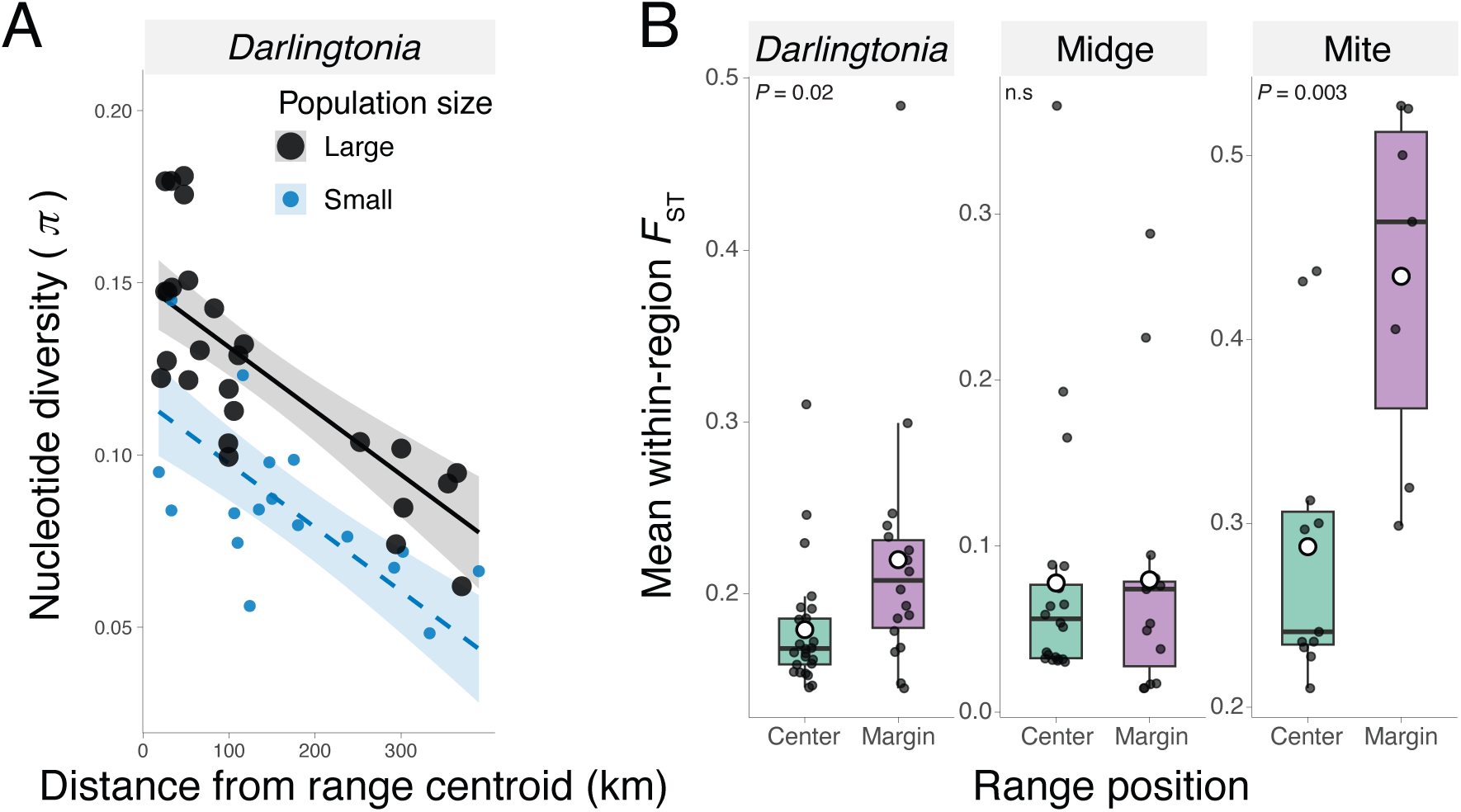
Tests of the central–marginal hypothesis. **(A)** Population-level nucleotide diversity (*π*) in the *Darlingtonia* nuclear dataset as a function of distance from the range centroid and population size. Lines show fitted linear regressions. **(B)** Mean within-region *F_ST_* for the *Darlingtonia* nuclear, *Metriocnemus edwardsi* (midge) COI, and *Sarraceniopus darlingtoniae* (mite) COI datasets in central and marginal portions of the range. Central populations comprise the Siskiyou and Trinity regions, and marginal populations the Oregon Coast and Sierra Nevada regions. Points represent individual populations, open circles indicate group means, and boxplots show the interquartile range. Reported *P* -values are from linear regressions comparing central and marginal populations, with n.s. denoting non-significant differences.

Marginal populations of *Darlingtonia* and *Sarraceniopus* were also more genetically isolated (**Fig. 3B**). Mean within-region *F_ST_*was significantly higher at the range margins than in central regions for the *Darlingtonia* nuclear dataset (*β* = 0.0407, *t*_42_ = 2.38, *P <* 0.05) and the mite dataset (*β* = 0.147, *t*_16_ = 3.53, *P <* 0.005), whereas the midge showed no central–marginal difference (*β* = 0.00186, *t*_35_ = 0.072, *P* = 0.94). Together, these results provide partial support for the central–marginal hypothesis for both *S. darlingtoniae* and its host plant, *D. californica*.

### Landscape resistance and effective migration

The top-performing MaxEnt model (feature classes = LQH, regularization multiplier = 1.5) showed good transferability to withheld data (continuous Boyce index = 0.85 ± 0.07). Model performance was generally similar across candidate feature-class and regularization settings, with validation AUC values of approximately 0.90, whereas the continuous Boyce index provided greater discrimination among candidate models and supported selection of the LQH model with a regularization multiplier of 1.5 (**Supplementary Figure S4**). Serpentine substrate was the most important predictor of habitat suitability (33.7%), followed by annual precipitation (17.1%), subsurface sand content (10.3%), temperature seasonality (8.5%), and isothermality (8.3%). The selected model predicted suitable habitat concentrated along serpentine corridors and coastal wetlands of northern California and Oregon, a pattern that closely matched the species’ known distribution (**Fig. 4A**).

**Figure 4:**
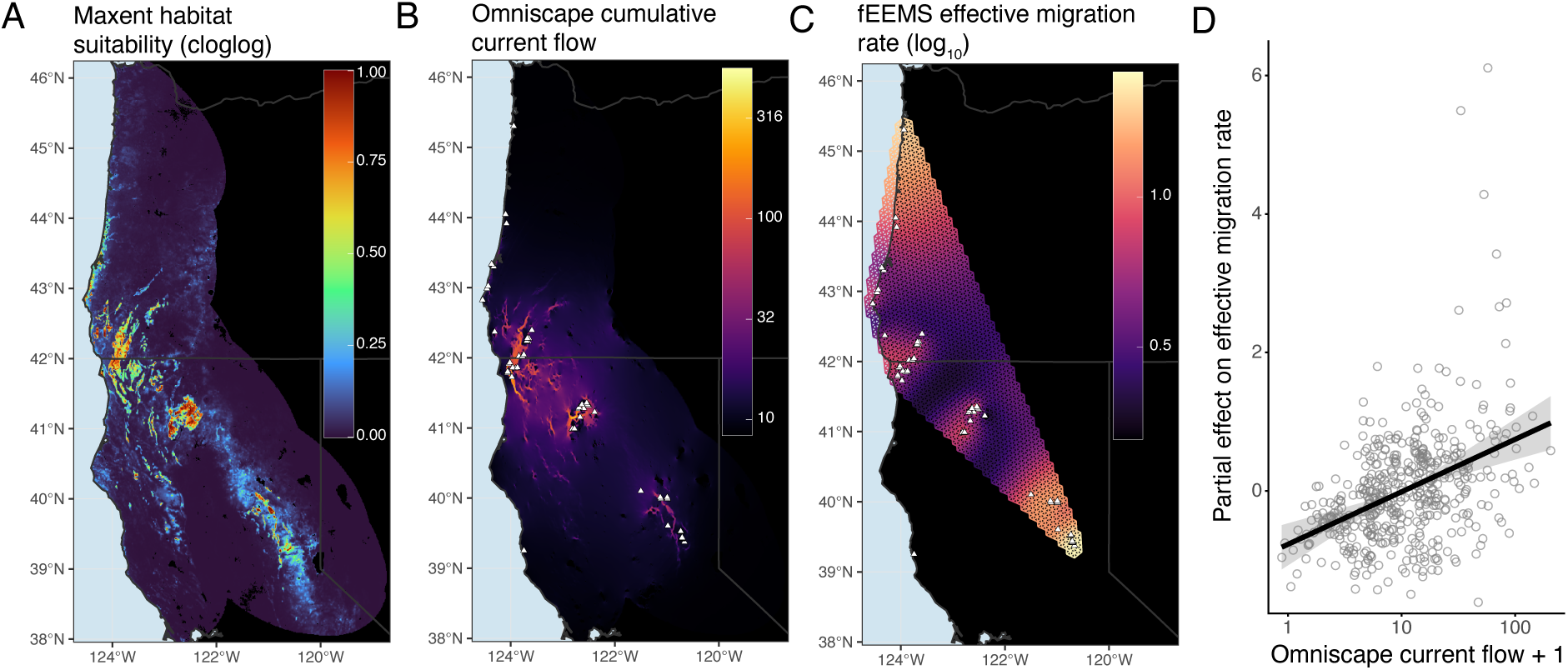
**(A)** MaxEnt-predicted habitat suitability for *D. californica*, with higher values indicating greater predicted suitability. **(B)** Cumulative current flow generated using Omniscape, with lighter areas indicating greater landscape connectivity among core habitat patches, defined as pixels with a resistance value of 1. **(C)** Effective migration rates inferred from *D. californica* genotypes using fEEMS, with darker colors indicating lower effective migration rates and therefore potential geographic barriers to gene flow. Sampling locations are indicated by white triangles. **(D)** Partial relationship between Omniscape cumulative current flow and fEEMS effective migration rate. The line shows the fitted effect of log_10_(current flow + 1) after accounting for spatial structure, the shaded band indicates the 95% confidence interval, and points show partial residuals.

Mapping cumulative current flow derived from Omniscape highlighted connective corridors within, but not among the four core regions (**Fig. 4B**). Independently, the fEEMS effective migration surface inferred from nuclear genotypes identified regions of reduced effective migration—candidate barriers to gene flow—between these same major regions (**Fig. 4C**).

As these two surfaces were derived from independent inputs, their spatial association provides an independent test of whether modeled connectivity predicts realized gene flow. In the GAM, Omniscape cumulative current flow was positively associated with fEEMS effective migration after accounting for spatial autocorrelation (*β* = 1.03, *t*_837.05_ = 6.33, *P <* 0.0001), and adding connectivity significantly improved fit over the spatial-only null (ΔAIC_null_ = 36.5) (**Fig. 4D**). Locations predicted to be more connective tended to show higher effective migration, indicating that the ENM-derived landscape identifies features that also facilitate gene flow.

### Isolation by distance and by resistance

The three organisms differed markedly in their isolation patterns (**Fig. 5**). *Darlingtonia* nuclear differentiation showed strong support for both isolation by distance (posterior prob[*β >* 0] = 1.00) and isolation by residualized resistance (prob[*β >* 0] = 1.00), indicating that geographic distance and landscape resistance each contribute independently to nuclear population structure. The midge *M. edwardsi* showed clear isolation by distance (prob[*β >* 0] = 1.00) but ambiguous resistance effects (prob[*β >* 0] = 0.37), while the mite *S. darlingtoniae* was positive for both (prob[*β >* 0] = 0.97 and 0.93) with wider credible intervals, as expected given its smaller number of sampled sites. Bayesian *R*^2^ indicated that geographic and residualized resistance distances together explained 32.6% of the variation in nuclear differentiation, 36.8% of the variation in midge differentiation, and 6.9% of the variation in mite differentiation. Including population-level multiple-membership effects increased the explained variation to 78.2%, 60.1%, and 67.5%, respectively, though the unique explanatory contributions of distance and resistance terms varied between 6% and 13% for the terms that lay significantly above zero (**Supplementary Table S4**).

**Figure 5:**
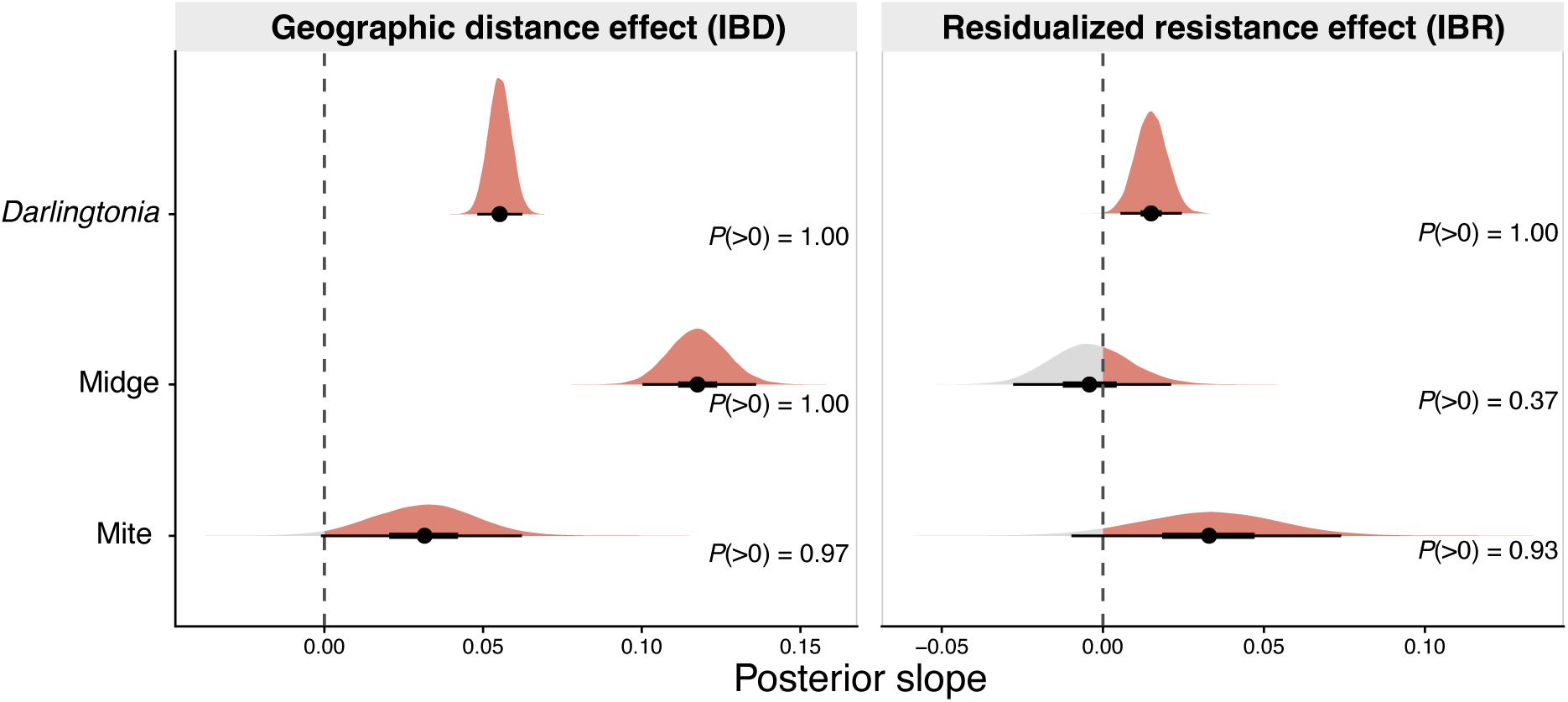
Posterior distributions of standardized slopes from Bayesian population-effects models relating pairwise *F_ST_* to geographic distance (isolation by distance) and residualized landscape resistance (isolation by resistance) for the *Darlingtonia* nuclear, *Metriocnemus edwardsi* (midge) COI, and *Sarraceniopus darlingtoniae* (mite) COI datasets. Horizontal intervals indicate the 50% and 95% credible intervals, and black points the posterior mean. Shaded portions of each posterior distribution indicate positive slope values. Values show the posterior probability of a positive slope, *P* (*β >* 0).

### Congruence of population structure across species

Pairwise Procrustes analyses of population *F_ST_* configurations indicated that each arthropod was spatially concordant with the host plant, but the two arthropods were not concordant with each other (**Supplementary Figure S5**). Host structure correlated significantly with both the midge (Procrustes *r* = 0.528, *P* = 0.0001) and the mite (*r* = 0.600, *P* = 0.0014), whereas the midge and mite configurations were not significantly congruent (*r* = 0.336, *P* = 0.285). Pairwise Mantel tests produced a similar pattern. *F_ST_*matrices were significantly correlated between host plant and midge (Mantel *r* = 0.443, *P* = 0.0001) and, more weakly, between host plant and mite (*r* = 0.316, *P* = 0.034), whereas the midge and mite matrices were not significantly correlated (*r* = 0.194, *P* = 0.137).

The three-way generalized Procrustes analysis, restricted to the 15 sites shared by all datasets, corroborated this picture (**Fig. 6**). The observed mean squared centroid deviation (0.024) was smaller than expected by chance when the plant configuration was permuted (*P* = 0.002), when the mite was permuted (*P* = 0.025), and when both arthropods were permuted jointly (*P* = 0.005), but not when the midge alone was permuted (*P* = 0.115). The shared congruence among the three organisms was therefore anchored by the host plant and, among the arthropods, driven more by the mite on this reduced site set. The midge’s weaker contribution, despite its significant pairwise concordance with the plant across its larger 37-site overlap, likely reflected the smaller, mite-limited set of sites available to the three-way test. Across both analyses, the two symbionts tracked the host plant but not each other, providing no evidence that the mite and midge are more tightly coupled to one another than to the plant.

**Figure 6:**
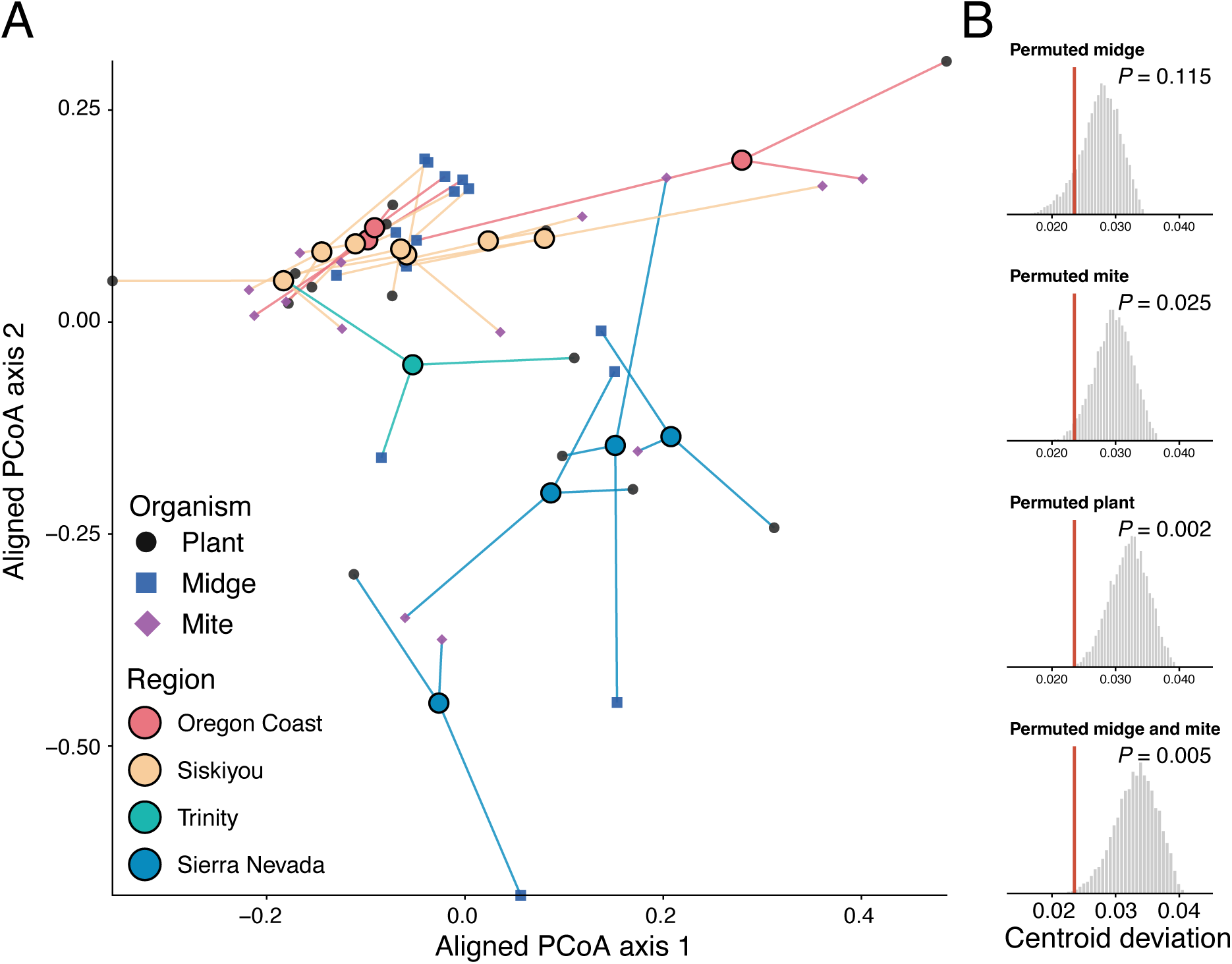
**(A)** Generalized Procrustes analysis of population configurations derived from the first two principal coordinate axes of nuclear plant, midge, and mite *F_ST_* matrices. For each locality, organism-specific scores are connected to their shared centroid; shorter connecting segments indicate greater agreement among species. Symbols denote organism and centroid colors denote geographic region. **(B)** Null distributions of mean squared centroid deviation generated by independently permuting locality identities for the midge, mite, plant, or both arthropods. Red lines indicate the observed three-way centroid deviation. *P* -values are the proportions of permutations with deviation less than or equal to the observed value.

## Discussion

We found that the pitcher plant *Darlingtonia californica* is strongly structured across its native range into four regional units, and that both genetic diversity and genetic isolation vary with position within that range. Populations near the range margins contained lower genetic diversity and were more differentiated from neighboring sites, in line with the central–marginal hypothesis. The mite *S. darlingtoniae* showed the same signature of elevated marginal isolation, whereas the midge *M. edwardsi* did not. Landscape connectivity predicted from an ecological niche model was positively associated with effective migration inferred independently from nuclear genotypes, indicating that modeled habitat structure captures features that also shape realized gene flow. Finally, both obligate arthropod symbionts showed isolation by distance and were spatially congruent with the host plant, yet not with one another.

### Regional structure, range margins, and the mating system

A recent RADseq survey of *Darlingtonia* across 14 sites in California recovered three genetic clusters, corresponding to the Six Rivers (Siskiyou), Shasta–Trinity, and Plumas (northern Sierra Nevada) National Forests, and interpreted this hierarchy as an outward radiation from a Klamath–Siskiyou refugium during a historical range expansion (Rice et al. 2026). That study was restricted to California and was based on a small panel of loci without individual genotype calls. Our range-wide sampling recovered the same three Californian clusters together with a fourth Oregon Coast cluster absent from that study, and found regional diversity highest in the Siskiyou region and lowest in the Sierra Nevada. We also independently recovered a Sierra Nevada origin for the introduced Albion Bog population.

Admixture among regions was limited. At *K* = 4, most *Darlingtonia* individuals were assigned to a single regional cluster, even where geographic barriers between neighboring regions (such as Oregon Coast and Siskiyou) are less clear. This pattern was consistent with the Omniscape surface, in which current flow was concentrated within rather than among regions, and with the fEEMS surface, which identified reduced effective migration near the same regional boundaries. The four regions therefore appear to represent stable, demographically differentiated populations. Gold Beach was an exception: its Siskiyou-like nuclear ancestry combined with an Oregon Coast plastid signature is consistent with historical seed exchange or chloroplast capture between these regions. Similar cytonuclear discordance has been documented elsewhere in Sarraceniaceae (Baldwin et al. 2023), although the low coverage and small number of plastid SNPs in our study warrant a cautious interpretation.

The central–marginal analyses showed that nuclear diversity declined with distance from the range center. Because the distance effect remained strong after population size was included, reduced diversity at the margins cannot be attributed solely to smaller local populations. Range position may instead reflect the effects of longer-term isolation, fewer neighboring populations, and a reduction in habitat suitability. Marginal populations were also more differentiated from neighboring populations within their regions. Together, the Oregon Coast and Sierra Nevada margins support the general prediction that populations near a species’ geographic limits experience reduced genetic connectivity (Vucetich and Waite 2003; Eckert et al. 2008).

Hierarchical AMOVA likewise documented strong structure at large and intermediate spatial scales. Differences among regions accounted for 27.4% of nuclear variation and differences among sites within regions a further 16.8%, placing 44.2% of the recovered variation above the individual-site level. Comparable regional partitioning has been reported in other pitcher plants: 16.9–28.5% of microsatellite variation occurred among population groupings in *Sarracenia alata* (Koopman and Carstens 2010), while regional structure in *S. purpurea* ranged from 3.8% within the western Lake Superior Basin (Karberg and Gale 2006) to 13.3% across an entire range-wide analysis (Karberg and Gale 2010). Although marker types and sampling designs differ, the strong regional component in *Darlingtonia* is consistent with its occurrence in isolated serpentine seeps and cold-water fens separated by extensive unsuitable terrain.

Variation among individuals within sites remained the largest single component, while relatively little variation occurred within individuals, consistent with the heterozygote deficits observed across all four regions. Clonality is unlikely to explain this pattern because every sampled ramet had a unique multilocus genotype. A more plausible explanation is the routinely documented self-pollination in *D. californica* (Meindl and Mesler 2011). Selfing reduces effective population size and increases the rate at which drift differentiates populations, potentially reinforcing the effects of habitat isolation (Wright et al. 2013).

The regional clusters are therefore most immediately relevant as conservation units rather than as evidence of cryptic species. Genetically differentiated lineages have been delimited within *S. alata* despite no apparent morphological differentiation (Carstens and Satler 2013; Zellmer et al. 2012), and geographic genetic units have been proposed as a conservation framework in *S. purpurea* (Karberg and Gale 2010). The four *Darlingtonia* regions experience distinct climates, and variation in thermotolerance, pigmentation, and stature has been noted across the species (McPherson and Schnell 2011). However, pending common-garden experiments and morphological comparisons, the regions are best regarded as strongly differentiated genetic units within a single species.

### Landscape connectivity and host gene flow

Nuclear differentiation in *D. californica* reflected independent contributions of geographic distance and landscape resistance. Distant populations were more differentiated, but populations separated by relatively unsuitable terrain were also more differentiated than geographic distance alone predicted. This result mirrors work in *S. alata*, where resistance distances based on major rivers explained genetic structure better than straight-line distance (Zellmer et al. 2012). In *Darlingtonia*, wide valleys lacking serpentine substrate and suitable hydrological conditions may form analogous barriers between occupied mountain systems.

Accounting for landscape resistance also helps explain why isolation by distance has often been weak or inconsistent in Sarraceniaceae. Geographic distance explained little differentiation in *S. alata* (Koopman and Carstens 2010) and *S. leucophylla* (Wang, Hamrick, et al. 2004), was detected in *S. oreophila* but not *S. jonesii* (Furches et al. 2013), and was absent in a range-wide analysis of *S. purpurea* (Karberg and Gale 2010). Our analysis identifies landscape resistance as one factor that can generate differentiation independently of distance, although substantial population-level variation remained after both predictors were included.

The positive association between Omniscape current flow and fEEMS effective migration provides a complementary spatial result. These surfaces were generated from independent environmental and genomic inputs, yet both identified high connectivity within regions and reduced connectivity near major regional boundaries. Their association persisted after spatial structure was included in the GAM, suggesting that the niche model identified landscape features that have also influenced realized gene flow. This is useful because landscape genetic studies commonly evaluate resistance models using pairwise genetic distances, whereas quantitative comparisons between independently estimated connectivity and migration surfaces remain rare (Spear et al. 2010).

This agreement should still be interpreted cautiously. Circuit-theoretic current flow represents the expected use of movement pathways under a random-walk model and is theoretically related to genetic coalescence and differentiation (McRae 2006; Fletcher Jr et al. 2022), but it is still not a direct measure of contemporary pollen or seed movement. Likewise, fEEMS estimates effective migration over genetic timescales rather than the immediate past. Our resistance surface also assumes a linear relationship between habitat suitability and movement cost (Keeley et al. 2016). Optimizing that relationship against the genetic data could improve fit (Peterman 2018), but would remove the statistical independence of the comparison. Future models that include pollinator and potential seed-disperser distributions may also provide a more mechanistic description of connectivity (Kling and Ackerly 2021; Davis et al. 2026).

### Dispersal mode and landscape responses

The three taxa diverged in ways that generally matched their dispersal biology. The host plant and mite showed greater differentiation at the range margins, whereas the midge did not. All three taxa exhibited isolation by distance effects, but only the host and mite showed evidence that landscape resistance explained additional variation beyond distance. The mite effect was less certain than that of the plant because fewer populations were sampled, but its direction was consistent with more restricted movement through unsuitable terrain. In contrast, the absence of a resistance effect in the winged midge suggests that the environmental features limiting plant and mite movement are comparatively less important to adult *M. edwardsi*.

Habitat configuration may still affect midge movement through mechanisms not represented by the plant-derived resistance surface. In *Metriocnemus knabi*, an inquiline of *S. purpurea*, bog size, host density, and patch connectivity explained up to 54% of variation in individual genetic distances, apparently through female oviposition behavior rather than matrix resistance (Rasic and Keyghobadi 2012a). Similarly, the pitcher-plant flesh fly *Fletcherimyia fletcheri* shows weak differentiation among bogs and isolation by distance only beyond approximately 10 km (Rasic and Keyghobadi 2012b). The landscape relevant to a flying pitcher-plant symbiont may instead more closely resemble a stepping-stone-like network of occupied breeding patches, whose connectivity depends more on patch adjacency than on the quality of the intervening matrix.

### Population genetic concordance among symbionts

Both arthropods showed significant spatial concordance with the host plant, yet no detectable concordance with each other. This pattern provides little support for frequent co-dispersal. Although laboratory observations suggest that mite deutonymphs can attach to adult midges, there is no evidence that phoresy is a common dispersal pathway (Fashing 2004). Moreover, midges leave pitchers before pupation (Jones 1916), limiting opportunities for mites to attach to newly emerged adults.

The magnitude of the mite–midge association was also consistent with independent host tracking. As a heuristic benchmark, if the two arthropods were related only through their separate associations with the host, their expected degree of correspondence would be the product of the two host–arthropod Procrustes correlations. This product was 0.317, close to the observed mite–midge correlation of 0.336. Although this is not a formal test of conditional independence, it provides no indication of additional correspondence attributable to shared dispersal. Congruence with the host can therefore arise through different mechanisms. Adult midges disperse actively, and their differentiation was dominated by geographic separation among host patches. Mites disperse locally as walking deutonymphs and perhaps occasionally through phoresy (Fashing 2004). Mites’ genetic differentiation was more sensitive to landscape resistance and more closely reflected constraints acting on the host. Therefore, shared population structure within an ecological community need not imply a common movement pathway.

This result extends an emerging view of pitcher-plant systems as evolutionary metacommunities. In the *S. alata* community, phylogeographic concordance with the host increased with ecological dependence. Obligate inquilines tracked host structure, whereas facultative spiders did not (Satler and Carstens 2019). Our results similarly indicate that obligate dependence can generate host–symbiont concordance. However, ecological dependence alone does not determine how that concordance arises and does not require obligate associates to be congruent with one another to maintain this highly specialized interaction.

Direct comparisons among taxa should account for differences in the underlying datasets. Plant structure was estimated from individually resolved nuclear genotypes, whereas arthropod structure was inferred from site-level mitochondrial COI sequences, and the smaller mite dataset reduced power to estimate tripartite concordance. Differences among taxa may also reflect the different modes of marker inheritance, effective population sizes, and sampling intensities as well as dispersal. Even so, the agreement among the central–marginal, IBD/IBR, and Procrustes analyses consistently suggests that that the winged midge *M. edwardsi* is less constrained by the host-derived resistance landscape than either *Darlingtonia* or its inquiline mite *S. darlingtoniae*.

### Conclusions

Our results contribute to an emerging consensus that even obligately interacting species need not share topologically congruent spatial genetic structure. Instead, they may converge on the same broad regional partition while responding to the landscape in different ways (Smith et al. 2019; Riley et al. 2023; Denis et al. 2026). Direct observations of dispersal in all three species would help resolve how the arthropod associates encounter and track their host plants, and potentially one another, across space.

In the applied context, these findings highlight the need to consider ecological interactions when using spatial genetic structure to guide management and conservation planning. This concern is especially relevant to serpentine montane fens, which are naturally sparse and sensitive habitats that support many rare and endemic plants (Jules et al. 2011). Changes in hydrology and fire regimes can rapidly eliminate suitable patches and reduce connectivity among those that remain.

Because species respond differently to this fragmented landscape, protecting habitat or connectivity for the host plant alone may not preserve the dispersal pathways required by its obligate associates. Nonetheless, their shared broad-scale genetic structure suggests that maintaining the regional network of fen habitats would benefit the metacommunity as a whole, particularly near range margins, where gains or losses of suitable habitat are most likely to reshape connectivity and gene flow (Anderson et al. 2009). The functional consequences of the reduced connectivity and genetic diversity observed near their range margins also remain unclear and warrant further investigation. More generally, extending landscape genetic analyses across multiple members of an interaction, rather than considering each species in isolation, will provide a more complete account of how dispersal sustains specialized ecological communities across space and time.

## Acknowledgments

We thank the Siletz Tribe, North Coast Land Conservancy, Oregon Parks and Recreation Dept., Siskiyou Field Institute, Oregon Dept. of State Lands, US Bureau of Land Management, US Forest Service, and the Hart family for permitting us access to *Darlingtonia* populations across Oregon and California. We also thank the individuals who contributed occurrence records, field assistance, laboratory support, and feedback on earlier versions of this work. Funding was provided by a Japan Cabinet Office Subsidy to OIST, and by a JSPS DC1 fellowship to N.H.

## Author Contributions

NH and DWA conceptualized the study, designed the research, and contributed to drafting the manuscript. NH, AEC, and DWA contributed to data collection. NH performed DNA extraction, sequencing, and sequence processing. NH led data analysis with contributions from DWA and MM. All authors commented on the manuscript and contributed to draft revisions.

