## Supplemental Methods, Tables, and Figures for "Connectivity and dispersal mode shape the landscape genetics of a carnivorous pitcher plant-arthropod metacommunity"

#### Supplementary contents

**1 Supplementary Methods** **2**
1.1 Plant DNA extraction, library preparation, and sequencing . . . . . . . . . . . . . . . . . 2
1.2 Organelle reference sequencing and assembly . . . . . . . . . . . . . . . . . . . . . . . . . 2
1.3 Arthropod DNA extraction, COI amplification, and sequencing . . . . . . . . . . . . . . . 2

**2 Supplementary Tables** **4**
Table S1: Environmental covariates used in ecological niche modelling . . . . . . . . . . . . . . 4
Table S2: ADAMIXTURE cross-validation across filtering schemes . . . . . . . . . . . . . . . . 5
Table S3: Within-region ADAMIXTURE cross-validation . . . . . . . . . . . . . . . . . . . . . 5
Table S4: Bayesian isolation-by-distance and resistance models . . . . . . . . . . . . . . . . . . 6

**3 Supplementary Figures** **7**
Figure S1: Geographic and residualized resistance distances . . . . . . . . . . . . . . . . . . . 7
Figure S2: fEEMS cross-validation . . . . . . . . . . . . . . . . . . . . . . . . . . . . . . . . . 8
Figure S3: Generalized Procrustes analysis workflow . . . . . . . . . . . . . . . . . . . . . . . 9
Figure S4: Ecological niche model selection . . . . . . . . . . . . . . . . . . . . . . . . . . . . 10
Figure S5: Pairwise Procrustes analyses . . . . . . . . . . . . . . . . . . . . . . . . . . . . . . 11
Figure S6: Chloroplast annotation for *Darlingtonia* . . . . . . . . . . . . . . . . . . . . . . . 12

**References** **13**

### 1 Supplementary Methods

#### 1.1 Plant DNA extraction, library preparation, and sequencing

Plant DNA was extracted from ground, dried leaf tissue using the MasterPure Complete DNA and RNA Purification Kit (Epicentre Technologies, Madison, WI). The volume of Proteinase K was increased from 1 to 5 $\mu$L, lysis was performed for 45 min at 1000 rpm and 56°C, and DNA was eluted in 25 $\mu$L TE buffer and stored at −20°C. DNA concentration was quantified using Qubit Broad Range and High Sensitivity dsDNA assays, and purity was estimated from A260/280 and A260/230 ratios measured with an Implen NanoPhotometer.

DNA was digested first with EcoRI-HF, bead purified, and then digested with BglII. Library quality was assessed for a random subset of samples before and after digestion using a Femto Pulse DNA electrophoresis system. ddRADseq libraries were prepared following Peterson et al. (2012), with adapters ligated using T4 DNA ligase, ligated fragments enriched using Dynabeads MyOne Streptavidin T1, and libraries amplified by PCR. Custom sequencing primers were used to reduce low sequence diversity around the restriction cut sites. Multiplexed libraries were sequenced on an Illumina NovaSeq 6000 SP flow cell over 300 cycles. In total, 199 DNA isolates were sequenced, with at least three individuals represented per collection site. Selected sites from each of the four principal geographic regions were represented by all eight sampled leaves.

#### 1.2 Organelle reference sequencing and assembly

Additional high-quality plant DNA extracts were selected for whole-genome shotgun sequencing on an Illumina platform using 151-bp paired-end reads. Raw-read quality was assessed using FastQC v0.11.9, and adapters and low-quality bases were removed with Trimmomatic v0.39 (Bolger et al. 2014). Organelle genomes were assembled *de novo* using NOVOPlasty v4.3.5 (Dierckxsens et al. 2017). For chloroplast assembly, *rbcL* was used as the seed and the *Actinidia arguta* plastome as a guiding reference. This produced three chloroplast contigs of 142,051, 12,733, and 12,885 bp, with an estimated mean organellar read depth of approximately 22,000×. The mitochondrial genome was assembled using NOVOPlasty's plant-mitochondrial mode, seeded with *cox1* and guided by the *Actinidia chinensis* mitochondrial genome, yielding a single 339,438-bp circular contig with an estimated mean depth of 2,383×. Assembly logs and overlapping termini were inspected to verify circularization, coverage consistency, and structural completeness. The chloroplast genome was then annotated using the GeSeq plastid annotation pipeline (Tillich et al. 2017) and its quality was verified based on the presence of known genes (Chang et al. 2026) and typical quadripartite structure (**Fig. S6**).

#### 1.3 Arthropod DNA extraction, COI amplification, and sequencing

DNA was isolated from preserved *Metriocnemus* larvae and *Sarraceniopus* mites using the MasterPure Complete DNA and RNA Purification Kit. Midges were flash-frozen in liquid nitrogen, ground with a micro-mortar and pestle, and extracted following the modified plant protocol. Mites were homogenized using a TissueRuptor (Qiagen, Inc.) and incubated in 600 $\mu$L lysis buffer for 2 h with shaking at 2000 rpm. Specimens were pooled by collection site because individual animals yielded insufficient DNA for amplification; consequently, the arthropod data do not support within-site, individual-level inference.

COI barcode amplicons were generated using Q5 Hot Start High-Fidelity 2X Master Mix (New England Biolabs, MA), 15–20 ng DNA, and each primer at 0.2 $\mu$M. Thermal cycling comprised 1 min at 95°C; 30 cycles of 10 s at 95°C, 30 s at 56°C, and 30 s at 72°C; and a final extension of 2 min at 72°C. We used the universal arthropod primer pair of Folmer et al. (1994): LCO1490, 5'-GGTCAACAAATCATAAAGATATTGG-3', and HCO2198, 5'-TAAACTTCAGGGTGACCAAAAAATCA-3', each appended to a Nextera transposase adapter. Amplification was verified by gel electrophoresis, and libraries were sequenced on an Illumina MiSeq using v3 chemistry over 600 cycles. Sequencing was successful for 57 midge and 45 mite samples.

#### 2 Supplementary Tables

Table S1: Environmental covariates considered when fitting the *Darlingtonia californica* ecological niche model. Collinear predictors were subsequently removed (see Methods); retained covariates are indicated in the final column.

| Name | Definition | Unit | Retained |
|---|---|---|---|
| *Bioclimatic* (Fick and Hijmans 2017) | | | |
| BIO1 | Mean annual temperature | °C | ✓ |
| BIO2 | Mean diurnal range | °C | |
| BIO3 | Isothermality (BIO2/BIO7 × 100) | – | ✓ |
| BIO4 | Temperature seasonality (SD × 100) | – | ✓ |
| BIO5 | Maximum temperature of warmest month | °C | ✓ |
| BIO6 | Minimum temperature of coldest month | °C | ✓ |
| BIO7 | Annual temperature range (BIO5 − BIO6) | °C | |
| BIO8 | Mean temperature of wettest quarter | °C | |
| BIO9 | Mean temperature of driest quarter | °C | |
| BIO10 | Mean temperature of warmest quarter | °C | |
| BIO11 | Mean temperature of coldest quarter | °C | |
| BIO12 | Annual precipitation | mm | ✓ |
| BIO13 | Precipitation of wettest month | mm | |
| BIO14 | Precipitation of driest month | mm | |
| BIO15 | Precipitation seasonality (coefficient of variation) | – | ✓ |
| BIO16 | Precipitation of wettest quarter | mm | |
| BIO17 | Precipitation of driest quarter | mm | ✓ |
| BIO18 | Precipitation of warmest quarter | mm | |
| BIO19 | Precipitation of coldest quarter | mm | |
| *Land cover* (Zanaga et al. 2021) | | | |
| Water | Area covered by water bodies >9 months per year | % pixel | ✓ |
| Wetland | Area dominated (>10%) by herbaceous wetland vegetation | % pixel | ✓ |
| Bare | Exposed soil, sand, or rock with <10% vegetation | % pixel | ✓ |
| Grassland | Area dominated (>10%) by grass and herbs | % pixel | ✓ |
| *Soil* (15 cm depth, mean) (Poggio et al. 2021) | | | |
| Soil cfvo | Volume fraction of coarse fragments >2 mm | % | ✓ |
| Soil nitrogen | Total nitrogen | g $kg^{-1}$ | ✓ |
| Soil pH | pH in $H_2O$ | – | |
| Soil sand | Sand (>0.05 mm) in fine earth | % | ✓ |
| Soil silt | Silt (0.002–0.05 mm) in fine earth | % | |
| Soil clay | Clay (<0.002 mm) in fine earth | % | |
| Soil soc | Organic carbon in fine earth | g $kg^{-1}$ | |
| *Substrate and hydrography* | | | |
| Serpentine | Ultramafic (serpentine) bedrock exposure | Binary | ✓ |
| SpringSeep | Euclidean distance to nearest mapped spring or seep | m | ✓ |

Table S2: Cross-validation (CV) error from `ADAMIXTURE` ancestry inference on *D. californica* nuclear SNPs, across combinations of minor allele frequency (MAF) and maximum locus missingness thresholds. Lower CV indicates better out-of-sample prediction of masked genotypes; the minimum for each filtering regime is shown in bold.

| MAF | Missingness | SNPs | $K$=2 | $K$=3 | $K$=4 | $K$=5 | $K$=6 | $K$=7 | $K$=8 | Best |
|---|---|---|---|---|---|---|---|---|---|---|
| 0.01 | 5% | 2,338 | 0.581 | 0.526 | **0.508** | 0.515 | 0.517 | 0.515 | 0.521 | 4 |
| 0.01 | 10% | 4,544 | 0.586 | 0.531 | **0.516** | 0.528 | 0.528 | 0.533 | 0.540 | 4 |
| 0.01 | 20% | 7,971 | 0.590 | 0.535 | **0.524** | 0.538 | 0.540 | 0.547 | 0.553 | 4 |
| 0.01 | 50% | 15,257 | 0.596 | 0.551 | **0.546** | 0.553 | 0.564 | 0.590 | 0.595 | 4 |
| 0.01 | 70% | 19,827 | 0.616 | 0.577 | **0.575** | 0.580 | 0.610 | 0.628 | 0.646 | 4 |
| 0.01 | 90% | 19,166 | 0.638 | **0.598** | 0.600 | 0.635 | 0.642 | 0.667 | 0.685 | 3 |
| 0.05 | 5% | 2,322 | 0.584 | 0.528 | **0.511** | 0.519 | 0.517 | 0.522 | 0.524 | 4 |
| 0.05 | 10% | 4,539 | 0.587 | 0.531 | **0.517** | 0.529 | 0.529 | 0.536 | 0.541 | 4 |
| 0.05 | 20% | 7,971 | 0.590 | 0.536 | 0.537 | **0.527** | 0.542 | 0.545 | 0.558 | 5 |
| 0.05 | 50% | 15,233 | 0.597 | **0.552** | 0.558 | 0.552 | 0.566 | 0.586 | 0.596 | 3 |
| 0.05 | 70% | 19,736 | 0.616 | 0.577 | **0.574** | 0.583 | 0.609 | 0.627 | 0.654 | 4 |
| 0.05 | 90% | 19,109 | 0.637 | **0.598** | 0.600 | 0.608 | 0.638 | 0.660 | 0.682 | 3 |

Table S3: Within-region cross-validation error from `ADAMIXTURE` ancestry inference on *D. californica* nuclear SNPs, following independent filtering and linkage-disequilibrium pruning of each regional subset. Values are means across five random seeds, and the minimum cross-validation error for each region is shown in bold. Cross-validation error was minimized at $K = 1$ in every region and in every individual run, indicating no detectable substructure within the four major ancestry clusters.

| Region | $n$ | SNPs | $K = 1$ | $K = 2$ | $K = 3$ | $K = 4$ |
|---|---|---|---|---|---|---|
| Oregon Coast | 27 | 752 | **1.193** | 1.300 | 1.562 | 1.785 |
| Siskiyou | 66 | 6414 | **0.906** | 0.942 | 1.026 | 1.105 |
| Trinity | 46 | 4555 | **1.016** | 1.049 | 1.126 | 1.204 |
| Sierra Nevada | 60 | 3158 | **0.862** | 0.887 | 0.949 | 1.014 |

Table S4: Posterior summaries and model-fit statistics for Bayesian MLPE models. Fixed-effect estimates and variance parameters are reported as posterior means with 95% credible intervals. Marginal Bayesian $R^2$ represents variation explained by geographic distance and residualized landscape resistance, whereas conditional Bayesian $R^2$ also includes population-level random effects. Predictor-specific contributions are expressed as the reduction in marginal Bayesian $R^2$ after excluding the focal predictor from the model.

| Dataset | Model quantity | Estimate | 95% credible interval | $P(\Delta R^2 > 0)$ |
|---|---|---|---|---|
| ***Darlingtonia*** | Intercept | 0.25 | 0.22–0.29 | — |
| | Geographic distance | 0.06 | 0.05–0.06 | — |
| | Residualized resistance | 0.01 | 0.01–0.02 | — |
| | Population-level SD | 0.11 | 0.08–0.13 | — |
| | Residual SD | 0.04 | 0.04–0.04 | — |
| | Marginal Bayesian $R^2$ | 32.6% | 27.3–37.3% | — |
| | Conditional Bayesian $R^2$ | 78.2% | 76.8–79.3% | — |
| | $\Delta R^2$: geographic distance | 12.6% | 6.7–18.1% | 1.00 |
| | $\Delta R^2$: residualized resistance | 7.3% | 1.4–12.9% | 0.99 |
| **Midge** | Intercept | 0.19 | 0.13–0.24 | — |
| | Geographic distance | 0.12 | 0.10–0.14 | — |
| | Residualized resistance | 0.00 | −0.03–0.02 | — |
| | Population-level SD | 0.16 | 0.12–0.21 | — |
| | Residual SD | 0.12 | 0.11–0.13 | — |
| | Marginal Bayesian $R^2$ | 36.8% | 30.9–42.2% | — |
| | Conditional Bayesian $R^2$ | 60.1% | 56.8–63.1% | — |
| | $\Delta R^2$: geographic distance | 8.0% | 1.3–14.5% | 0.99 |
| | $\Delta R^2$: residualized resistance | 0.0% | 0.0–6.2% | 0.43 |
| **Mite** | Intercept | 0.35 | 0.27–0.43 | — |
| | Geographic distance | 0.03 | 0.00–0.06 | — |
| | Residualized resistance | 0.03 | −0.01–0.07 | — |
| | Population-level SD | 0.16 | 0.11–0.24 | — |
| | Residual SD | 0.08 | 0.07–0.09 | — |
| | Marginal Bayesian $R^2$ | 6.9% | 0.2–20.0% | — |
| | Conditional Bayesian $R^2$ | 67.5% | 61.1–72.3% | — |
| | $\Delta R^2$: geographic distance | 6.5% | 0.0–19.7% | 0.96 |
| | $\Delta R^2$: residualized resistance | 6.0% | 0.0–19.2% | 0.90 |

*Notes:* Geographic distance and residualized resistance were standardized before model fitting. Population-level SD is the standard deviation of the random intercept for the two populations represented in each pairwise comparison. The predictor-specific $\Delta R^2$ is the difference between the marginal Bayesian $R^2$ of the full model and that of a reduced model excluding the corresponding predictor, and represents change in explained variation following predictor exclusion.

### 3 Supplementary Figures

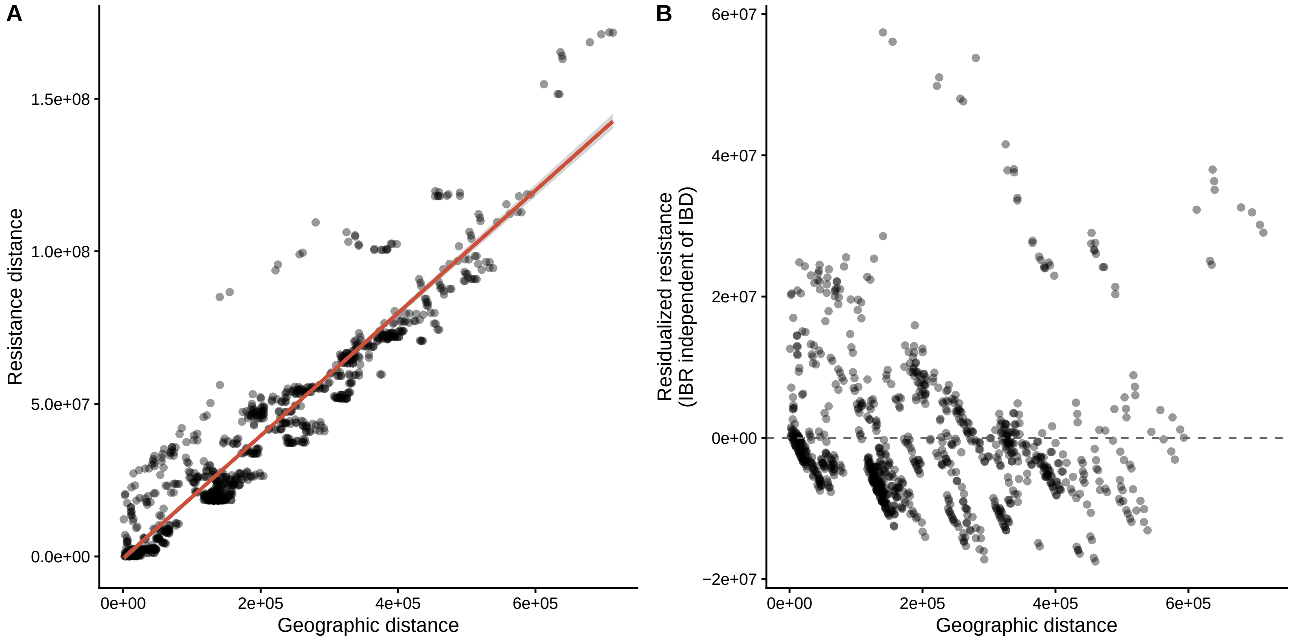


Figure S1: **(A)** Relationship between pairwise geographic distances and pairwise resistance distances for *D. californica*. **(B)** Relationship between pairwise geographic distances and residualized resistance distances for use in MLPE models.

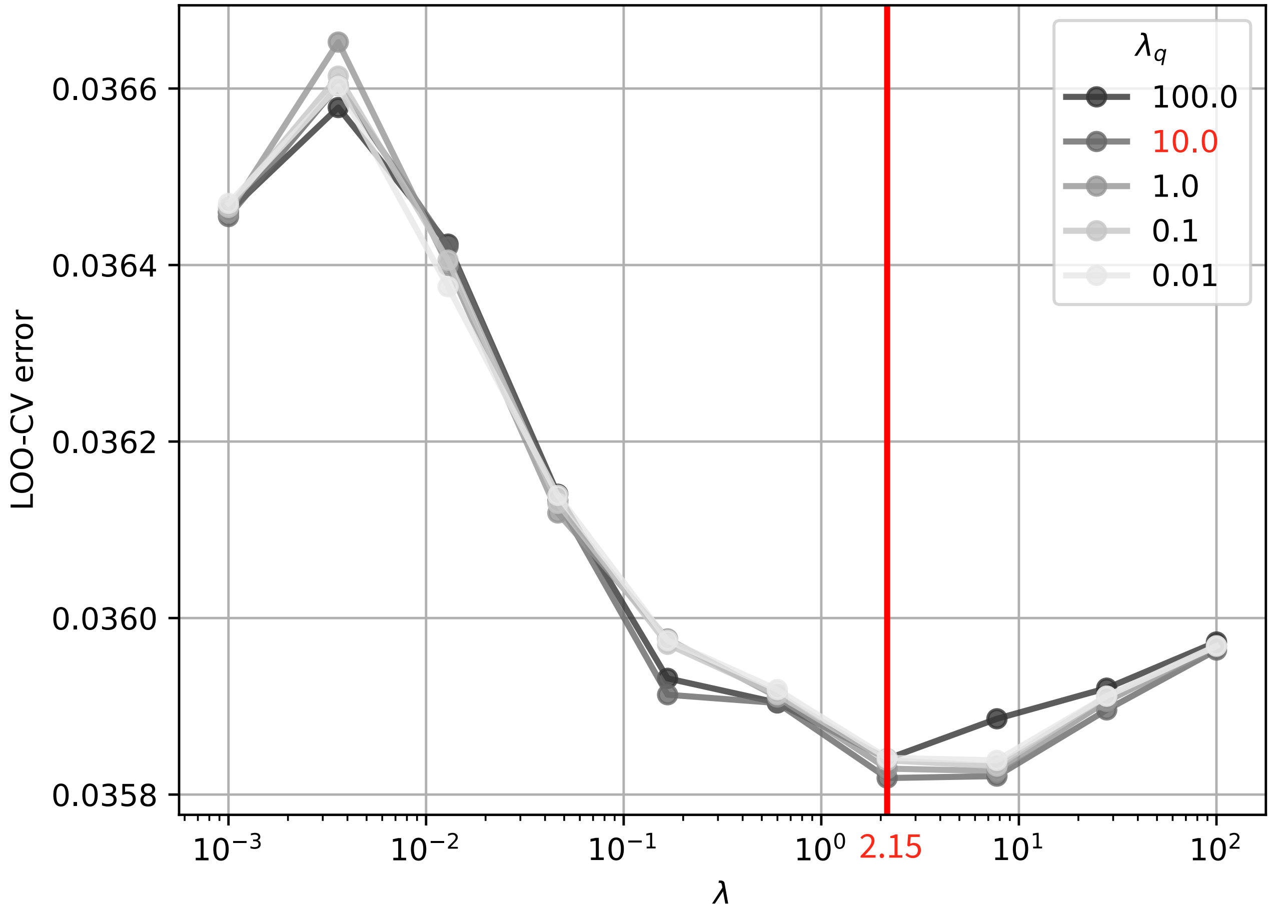


Figure S2: Leave-one-out cross-validation (LOO-CV) error used to select the regularization parameters for the fEEMS model. Cross-validation error is shown across values of the spatial regularization parameter $\lambda$ for five values of the residual-variance regularization parameter $\lambda_q$. The red numbers show the parameter combination yielding the minimum LOO-CV error.

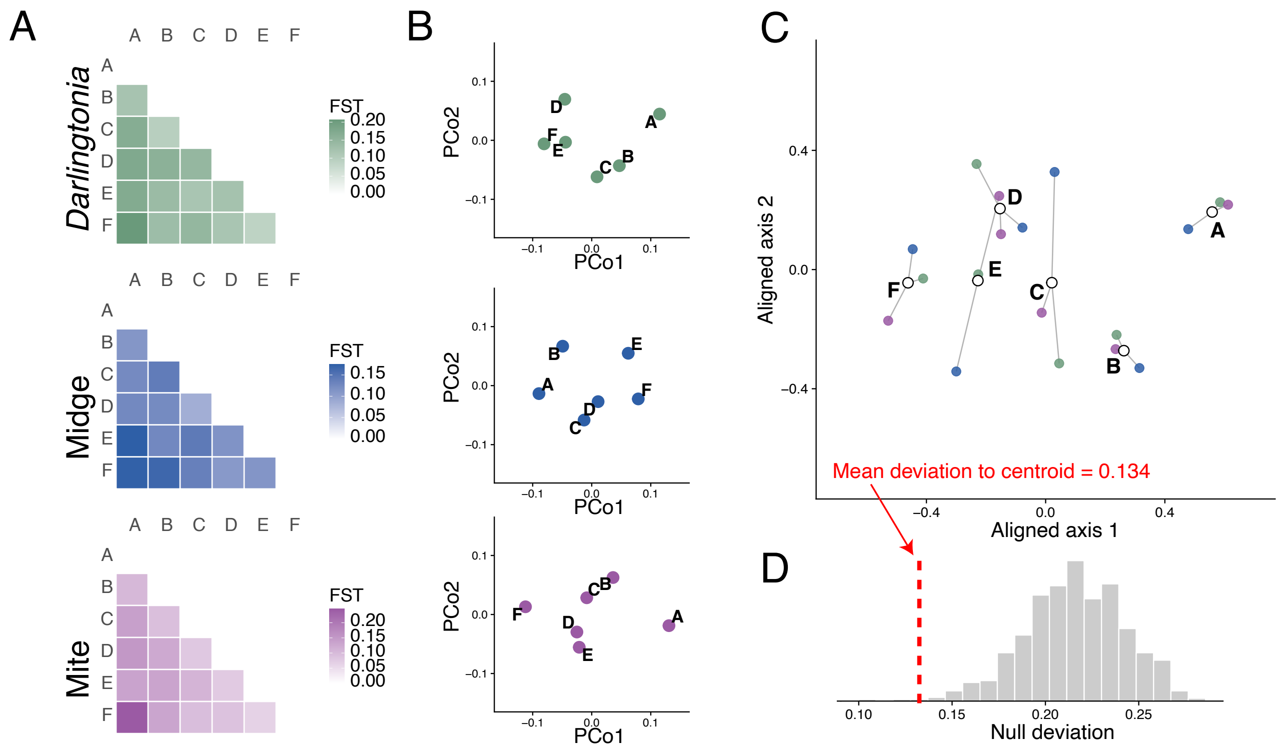


Figure S3: Overview of the generalized Procrustes analysis used to measure concordance among population genetic structures. **(A)** Pairwise $F_{ST}$ matrices are calculated independently for each species. **(B)** Each distance matrix is ordinated using principal coordinate analysis (PCoA), producing an independent representation of population genetic relationships. **(C)** These configurations are aligned by generalized Procrustes analysis, which translates, rotates, and scales the ordinations to minimize the mean Euclidean deviation of corresponding populations from their shared centroid. The mean deviation across all populations provides a measure of concordance among species. **(D)** Statistical significance is assessed by comparing the observed mean deviation with a null distribution generated by repeatedly permuting the population correspondence for one or more datasets prior to Procrustes alignment. The proportion of permutations producing an equal or larger mean deviation gives the permutation $P$-value.

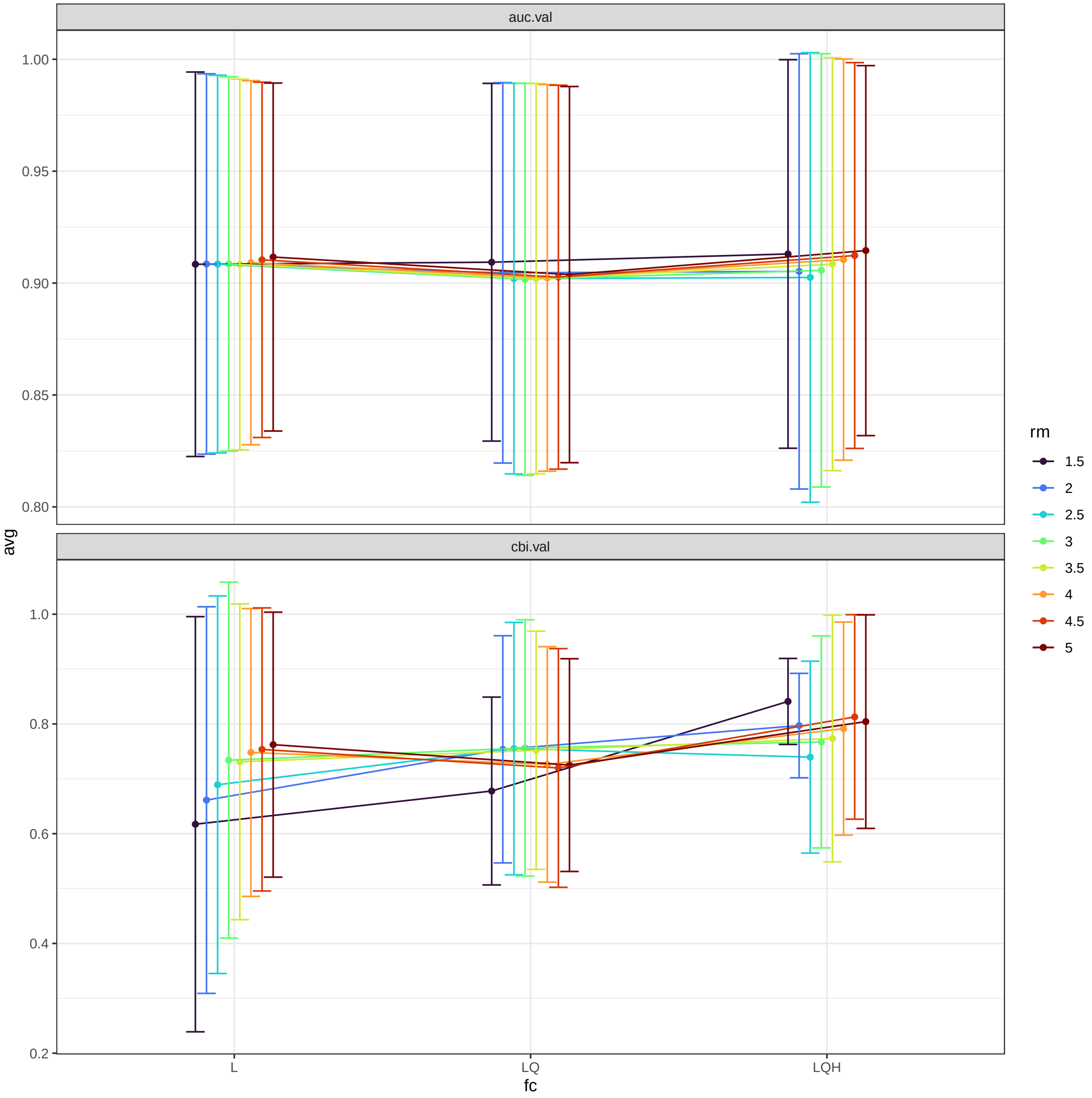


Figure S4: Cross-validation performance across feature-class (`fc`) and regularization-multiplier (`rm`) combinations for MaxEnt models. Points show the mean test area under the receiver operating characteristic curve (`auc.val`; upper panel) and continuous Boyce index (`cbi.val`; lower panel), with error bars indicating variation among spatial partitions. Feature classes were linear (`L`), linear–quadratic (`LQ`), and linear–quadratic–hinge (`LQH`); colors show regularization multipliers ranging from 1.5 to 5.

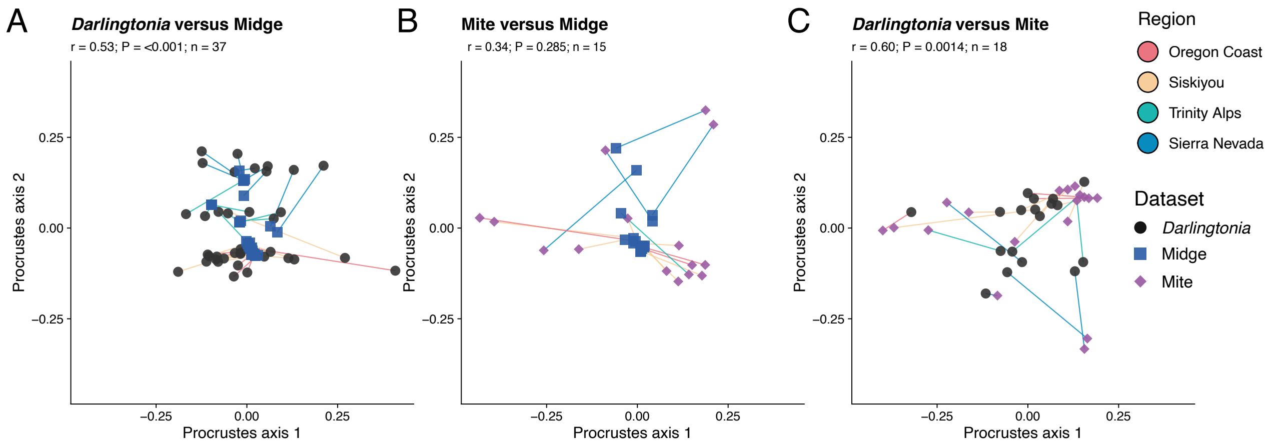


Figure S5: Pairwise Procrustes analyses for **(A)** *Darlingtonia* nuclear DNA and *M. edwardsi* COI barcode data. **(B)** *M. edwardsi* and *S. darlingtoniae* COI barcode data. **(C)** *Darlingtonia* nuclear DNA and *S. darlingtoniae* COI barcode data. *P*-values indicate the probability of obtaining an equal or smaller mean Procrustes deviation under the null hypothesis of no correspondence between datasets; small *P*-values therefore indicate significantly stronger concordance than expected by chance. Lines are colored by region. Point shapes indicate species.

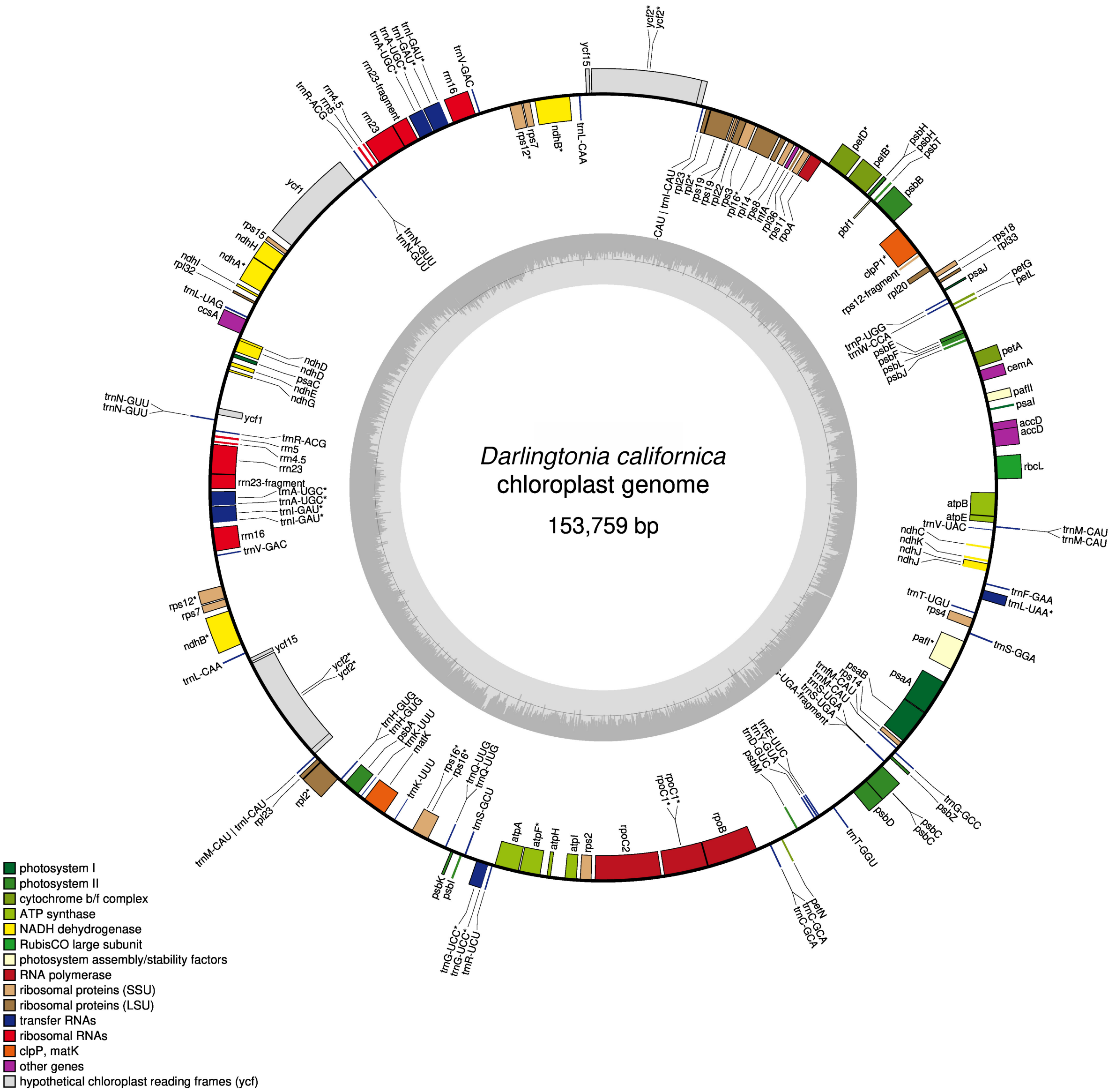


Figure S6: Circular map of the *Darlingtonia californica* plastome. Genes drawn outside or inside the circle are transcribed counter-clockwise or clockwise, respectively. Colors denote functional categories. Asterisks mark intron-containing genes. The inner grey ring shows GC content in 200 bp windows, with a midline at 50%. Genes of the NADH dehydrogenase (*ndh*) complex are extensively pseudogenized. *ndhE* and *ndhH* retain intact reading frames, while the remaining *ndh* genes are disrupted by frameshifts and premature stop codons. *ndhF* could not be annotated and is shown as an unannotated region.
